# Development of NanoBRET cellular target engagement assays in primary neurons for activating mutants of p21-activated kinase 1

**DOI:** 10.64898/2026.05.03.722513

**Authors:** Jacob L. Capener, Abdiel Badillo-Martinez, Bassel Awada, Zachary W Davis-Gilbert, Thomas W. Kramer, Cameron S. Blair, Frances M. Bashore, Hassan Al-Ali, Alison D. Axtman

**Affiliations:** Structural Genomics Consortium, UNC Eshelman School of Pharmacy, University of North Carolina at Chapel Hill, Chapel Hill, North Carolina 27599, USA; Department of Biochemistry and Molecular Biology, University of Miami Miller School of Medicine, Miami, FL, USA; Miami Project to Cure Paralysis, Department of Neurological Surgery, University of Miami, Miami, FL, USA; Department of Neurological Surgery, Peggy and Harold Katz Drug Discovery Center, Sylvester Cancer Comprehensive Center, and Frost Institute for Data Science and Computing, University of Miami, Miami, FL, USA

## Abstract

The p21-activated kinases (PAKs) are a group of serine-threonine kinases central to multiple signaling pathways that govern cell survival and proliferation. Aberrant activity of PAK1, the most well characterized member of the PAK family, drives progression of several malignancies and brain disorders, including Alzheimer’s disease and neurodevelopmental disorders. Despite growing interest in PAK1 as a drug target for these diseases, there is no assay to evaluate the intracellular target engagement of PAK1 inhibitors. To address this need, we developed first-in-class NanoBRET assays for wild-type PAK1 and a neurodevelopmental disorder-causing gain-of-function PAK1 mutant. Furthermore, we executed our novel PAK1 NanoBRET assay to evaluate target engagement of PAK1 inhibitors in primary hippocampal neurons. To the best of our knowledge, this is the first demonstration of a NanoBRET cellular target engagement assay in primary neurons, thereby increasing the relevance of our work by confirming PAK1 inhibitor binding to the aberrant form of the protein in primary neurons.

## Introduction

The group 1 p21-activated kinases (PAKs) are a class of conserved proteins with high sequence homology and roles in controlling cytoskeletal motility, cell survival, and proliferation^1^. The PAK kinases have overlapping and divergent functions. Among the three group 1 PAKs (PAK1–3), PAK1 has emerged as the most sought-after drug target because it regulates key signaling pathways, including PI3K/AKT, MAPK, and WNT/β-catenin^1,2,3,4^. PAK1 promotes cancer cell survival in hypoxic conditions and accelerates angiogenesis, contributing to cancer metastasis^1,5^. Additionally, activating mutations in proteins upstream of PAK1, such as Rac1 or CDC42, or PAK1 gene amplification, are implicated in pancreatic, hepatocellular, breast, and prostate cancer^1,2,6,7^. Outside of cancer, recent reports tie PAK1 signaling pathways to Alzheimer’s disease (AD) and Fragile X syndrome^8,9^. Furthermore, aberrant PAK1 activity resulting from specific single point mutations in PAK1 has been shown to cause neurodevelopmental disorders^10,11^. Specifically, in one example identified in 2018, two distinct forms of PAK1 with different activating mutations, Y131C and Y429C, were separately identified in patients with developmental delays^11^. Mechanistically, these mutations disrupt the autoinhibitory regulation of PAK1 (Figure 1A). Historically, structural studies largely converged on the hypothesis that group 1 PAK members are regulated through a *trans*-inhibited dimer conformation in which the N-terminal autoinhibitory domain (AID) of one PAK monomer binds to and inhibits the C-terminal catalytic domain of a second PAK monomer^12^. Recent structural work examining PAK1 and full-length PAK2, however, has demonstrated that a *cis* monomer autoinhibition mechanism is favored for the group 1 PAKs^13,14^. This autoinhibition mechanism is essential for maintaining normal PAK activity in the cell. Critically, when this autoinhibition process is disrupted by these activating mutations, constitutive PAK1 activity drives disease-causing phenotypes.

**Figure 1.**
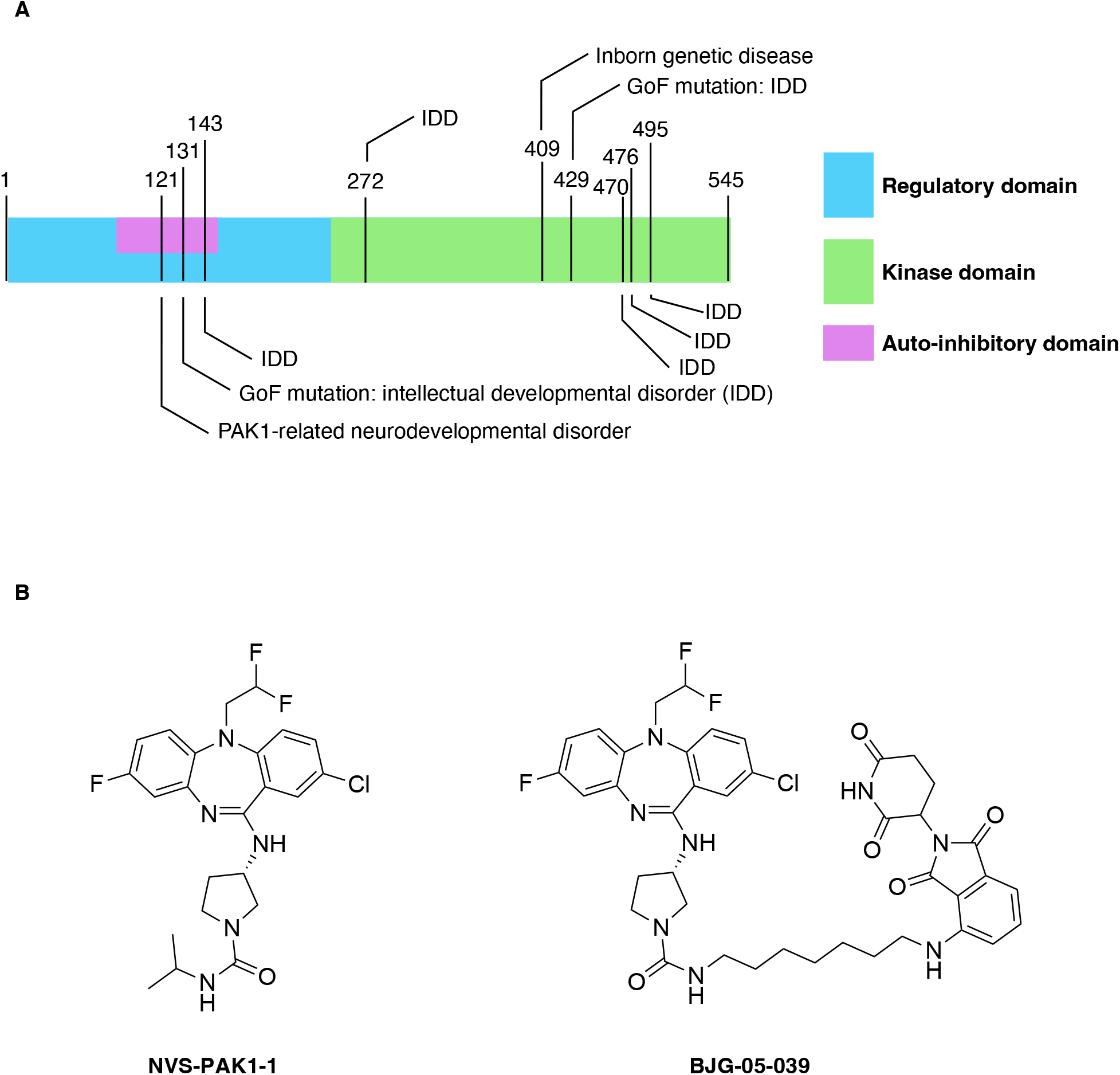
Diagram of PAK1 structure and available chemical tools. (a) Schematic displaying various domains and known disease-associated mutations on PAK1, including the gain-of-function mutations at residues 131 and 429. (b) Chemical structure of the Novartis PAK1 probe, NVS-PAK1-1, and the PAK1 PROTAC, BJG-05-039.

To combat diseases proposed to be driven by aberrant PAK1 activity, several groups have developed PAK1 inhibitors that have been evaluated in both cellular and in vivo contexts^7^. PAK1 inhibitors have demonstrated significant promise for several cancers and, more recently, AD^15^. One inhibitor, PF-3758309, was even advanced to phase 1 clinical trials for the treatment of patients with advanced solid tumors before failing due to poor bioavailability. Later, it was determined that pan-group 1 PAK inhibitors, such as PF-3758309, result in cardiotoxicity due to PAK2 inhibition^16^. Currently, no orthosteric inhibitors have achieved selectivity within the highly homologous group 1 PAKs. Thus, all orthosteric PAK1 inhibitors have the liability of PAK2 inhibition. As an alternative strategy, Novartis developed an allosteric, ATP-competitive dibenzodiazepine compound, NVS-PAK1-1, that potently inhibited PAK1, with excellent kinome-wide selectivity^17^. Notably, NVS-PAK1-1 lacks binding affinity for other PAK isoforms, including PAK2. Thus, NVS-PAK1-1 does not elicit the cardiotoxicity observed with other PAK1 inhibitors^15^. NVS-PAK1-1 binds to an allosteric pocket adjacent to the ATP-binding site of PAK1, conferring its selectivity, and interferes with ATP binding. This is the only PAK1 inhibitor that meets the chemical probe criteria set by the Structural Genomics Consortium (SGC)^18^. Use of NVS-PAK1-1 in cellular studies has revealed distinct functions of PAK1 and identified functional redundancies among the group 1 PAK isoforms^15,19,20^. Furthermore, a degrader derived from NVS-PAK1-1, BJG-05-039, has been used alongside NVS-PAK1-1 to dissect the scaffolding functions of PAK1^21^. Although promising data has been generated using PAK1 inhibitors and degraders, no molecules have yet demonstrated the requisite properties to support clinical trials beyond the aforementioned phase 1 study.

Despite the need for new PAK1 inhibitors, there are currently no methods to evaluate the cellular target engagement of a compound to PAK1. Currently, the gold standard for assessing cellular target engagement is using Nanoluciferase (NLuc)-based bioluminescence resonance energy transfer (NanoBRET)^22,23^. NanoBRET measures the displacement of a bifunctional tracer molecule, known as a BRET probe, that contains an inhibitor moiety and a NanoBRET 590 dye. The BRET probe gets excited in the proximity of NLuc, producing a BRET signal. The binding of a test compound to the same site on an NLuc-tagged protein as the BRET probe results in the displacement of the tracer and a quantifiable loss of the BRET signal. Currently, all methods for assessing the inhibitory potency of PAK1-targeting compounds rely on biochemical or substrate phosphorylation assays. These assays report on kinase activity rather than the affinity of a small molecule for the kinase. Biochemical assays, while useful, do not account for variables that may influence in-cell binding, such as compound cell permeability, post-translational modifications of PAK1, or the intracellular concentration of ATP that results in binding competition for most PAK1 inhibitors^24^. While many of these variables are accounted for when using in-cell substrate phosphorylation assays, these methods are indirect and may be influenced by compensatory signaling or other off-target interactions^23^. The NanoBRET cellular target engagement assay complements other assay formats by providing direct, quantitative confirmation of binding to the target of interest through tracer displacement within a living cell. The absence of a cellular target engagement assay for PAK1 represents a significant obstacle to the development and optimization of cell-active PAK1 inhibitors.

In this work, we addressed the need for a cellular target engagement assay for both wild-type and gain-of-function PAK1 mutants by enabling NanoBRET assays for both forms. This work required the development of a cell-permeable fluorescent BRET probe, which was derived from NVS-PAK1-1. Using our newly developed assays, we evaluated the cellular affinity of several reported PAK1 inhibitors for both wild-type PAK1 and disease-causing PAK1 mutants. Finally, we ported the assay to primary hippocampal neurons to assess the affinity of these inhibitors for PAK1 in a target and highly disease-relevant cell model. To our knowledge, this represents the first NanoBRET cellular target engagement assays performed in primary neurons, and it also demonstrates the robustness of the assay in human and non-human (rat) cells.

## Results and Discussion

### BRET Probe Development

The development of a BRET probe is a significant obstacle to establishing a NanoBRET assay for a protein target. To achieve a BRET signal, a NanoBRET 590 dye must be covalently linked to a known ligand at a site that does not perturb binding to the target protein. Additionally, this bifunctional BRET probe must remain cell-permeable to be used in a live-cell assay. For this reason, the historical strategy for kinase NanoBRET assays has been to employ promiscuous BRET probes to enable multiple assays using a single tracer molecule^22^. However, for many kinases that are not bound by existing promiscuous ligands, tracers have been synthesized from more selective ligands to establish a NanoBRET assay^25,26,27^. In the case of PAK1, NVS-PAK1-1 is among the most potent and best studied compounds. Thus, we decided to generate our BRET probe through chemical modification of NVS-PAK1-1.

As part of our tracer design strategy, we modified NVS-PAK1-1 at the solvent-exposed isopropyl urea site, which was previously exploited in the development of the PAK1 degrader, BJG-05-039 (Figure 1B)^21^. To couple NVS-PAK1-1 to the NanoBRET 590 dye, we used several alkyl or polyethylene glycol (PEG) linkers of varying lengths to generate four distinct BRET probes (Scheme 1 and Figure 2A). We then evaluated all four tracers against the full-length PAK1 protein tagged with a C-terminal NLuc, and, interestingly, no BRET signal was observed (Figure 2B). As our BRET probes varied in linker lengths and chemical composition, and we included the octyl linker used for BJG-05-039, which retained binding to PAK1, we concluded that it was unlikely that we had perturbed binding through linker attachment. When we had previously developed a SYK NanoBRET assay, a suite of BRET probes with varying linker lengths retained binding to the wild-type (WT) form of SYK but failed to produce a NanoBRET signal in intact and permeabilized cells. For SYK, BRET probes only produced a signal through imparting gain-of-function SYK mutations that disrupted the binding of the AID to the ATP-binding site, which physically prevented tracers from binding^25^. It was proposed that a similar strategy may be a viable approach for PAK1.

**Scheme 1.**
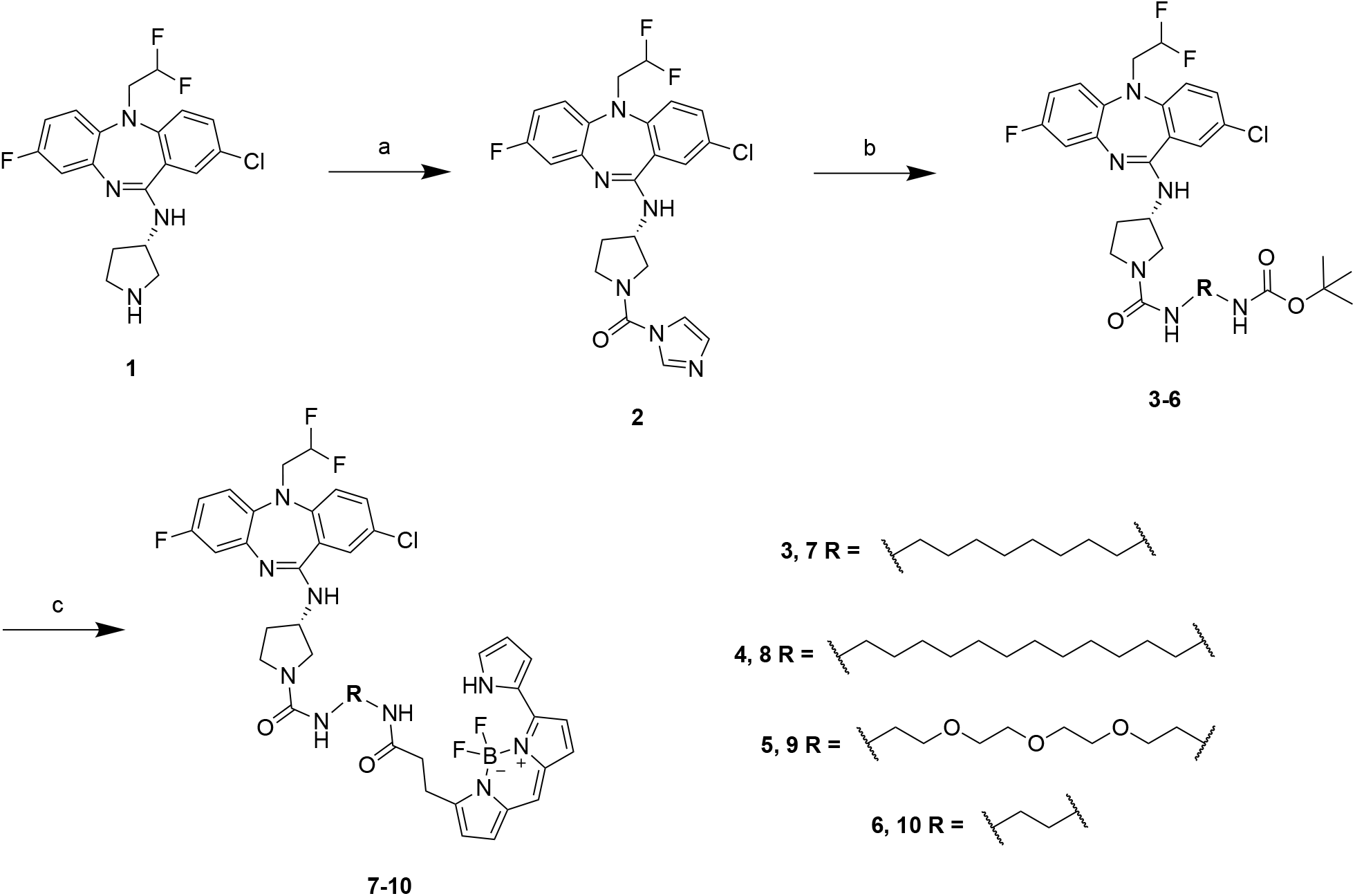
Synthesis of Tracers 7-10. Reagents and conditions: (a) CDI, TEA, DCM, 0°C to r.t., 24 h, 77% yield. (b) 1) MeI, ACN, r.t., 24 h. 2) HATU, DIPEA, Amine-R, DMF, r.t., 24 h, 3–44% yield over two steps. (c) 1) DCM, TFA, r.t., 4 h. 2) 2,5-dioxopyrrolidin-1-yl 3-(5,5-difluoro-7-(1*H*-pyrrol-2-yl)-5*H*-5λ^4^,6λ^4^-dipyrrolo[1,2-*c*:2′,1′-*f*][1,3,2]diazaborinin-3-yl)propanoate, DIPEA, DMF, r.t., 24 h, 30–80% yield over two steps

**Figure 2.**
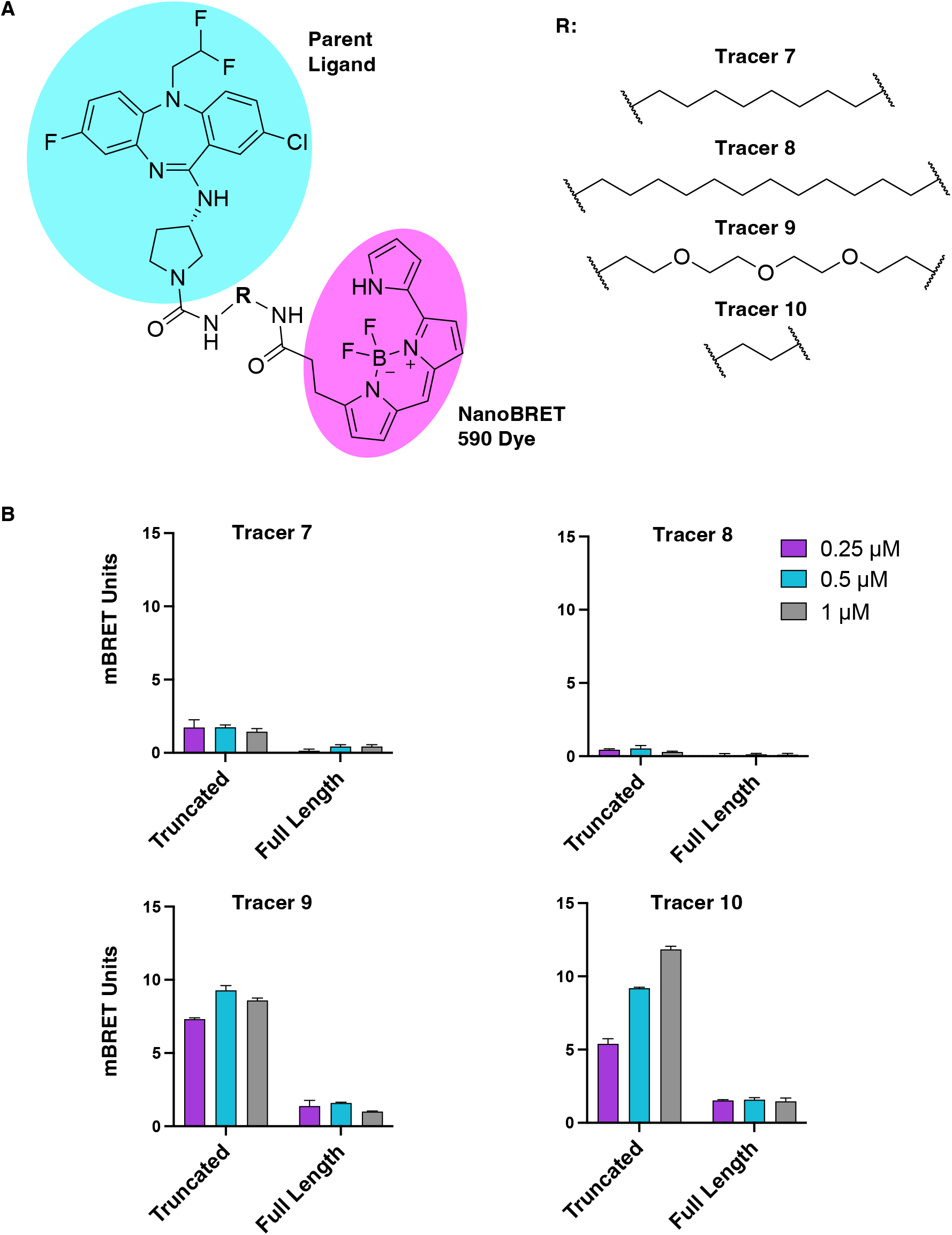
BRET probe linker optimization and resulting BRET signal. (a) Structure of our NanoBRET tracer with NVS-PAK1-1 highlighted in blue, NanoBRET 590 dye highlighted in purple, and the varied linker structures shown on the right with the corresponding tracer name. (b) BRET signal elicited by each tracer at 0.25, 0.5, and 1 μM for both the full-length and truncated WT PAK1 constructs. (n=1, performed in technical duplicate)

### Enabling NanoBRET Assays for Different PAK1 Variants

To test whether AID binding to the ATP-binding site precluded tracer engagement, we screened our BRET probes against a truncated form of PAK1 that excluded the AID (248–545) region and bore an C-terminal NLuc tag. When using this truncated PAK construct, tracers **9** and **10** produced a dose-dependent NanoBRET signal greater than two-fold over background, which was competitive with NVS-PAK1-1 (Figure 2B). The signal produced by both tracers was sufficient for NanoBRET assay development, and we elected to advance tracer **10** to further assay establishment experiments using the truncated form of the kinase. To this end, we determined the EC_50_ of the tracer for truncated wild-type PAK1 to be 310 nM. Using the tracer EC_50,_ a K_i,app_ value can be calculated via the Cheng-Prusoff equation (Figure 3A). Additionally, consistent with the competitive binding model captured by the Cheng-Prusoff equation, we performed a competition experiment with tracer **10** and observed a linear relationship between the NVS-PAK1-1 IC_50_ value and tracer concentration (Figure 3B). While this assay that employs truncated PAK1 represents the first model for cellular assessment of target engagement for PAK1, previous work has shown that truncated kinase domains may not capture the behavior of the native enzyme^24^. Specifically, assays with isolated kinase domains may yield different K_m_ values for ATP and divergent binding affinities for inhibitors that exhibit conformational selectivity compared to the full-length protein^28^. Thus, full-length proteins better represent the endogenous state of the protein.

**Figure 3.**
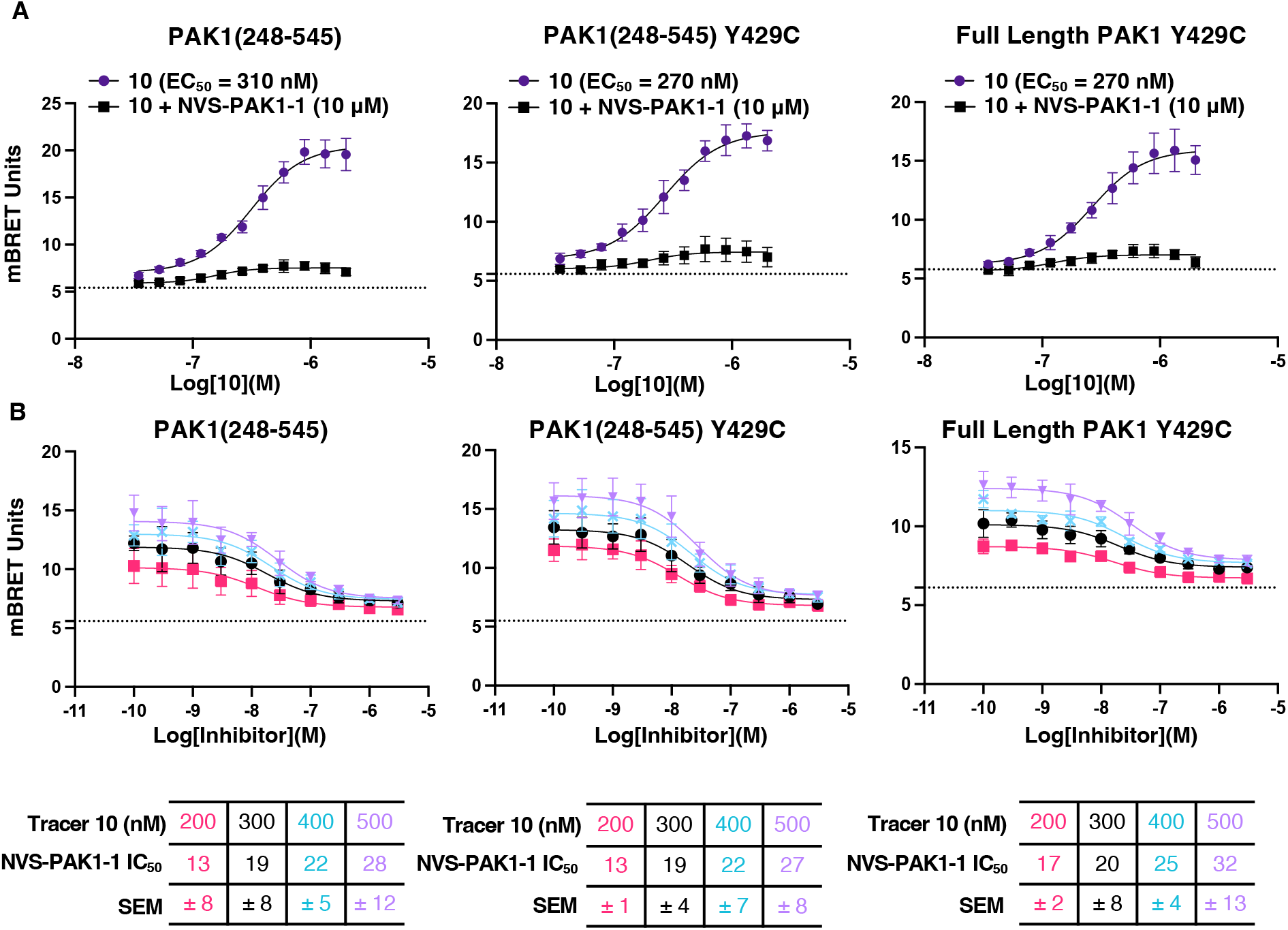
Development of the NanoBRET assay for PAK1(248–545), PAK1(248–545) Y429C, and full-length PAK1 Y429C. (a) Tracer titration experiments for tracer EC_50_ determination with an 11-point tracer titration study for tracer **10** with and without NVS-PAK1-1. (b) Competition experiments with varying doses of tracer **10** and IC_50_ determination of NVS-PAK1-1 at each tracer concentration. The table reports the NVS-PAK1-1 IC_50_ at each tracer concentration, along with the standard error of the mean (SEM). The data are not background-subtracted, and the horizontal dotted line indicates the background level. All data is reported as n=3 ± SEM as indicated.

To develop an assay for the full-length form of PAK1, we used disease-causing gain-of-function PAK1 mutations that disrupt AID binding due to a single point mutation. We selected a well-characterized mutation at tyrosine 429 of PAK1, which has been shown to cause a rare neurodevelopmental disorder^11^. Previous studies have demonstrated that the pathogenic mechanism linked to this mutation is constitutive PAK1 activity, resulting from a significant reduction in AID binding. Specifically, the Y429C mutant decreased AID binding by 70% compared to WT PAK1, making this form of PAK1 an excellent candidate for NanoBRET assay development^11^. First, to ensure that the mutation at tyrosine 429 did not alter inhibitor or tracer binding, we developed an assay using the truncated form of PAK1 (248–545) harboring the Y429C point mutation. In both the tracer titration and the competition experiments using tracer **10**, the mutated and wild-type truncated constructs exhibited similar assay windows and IC_50_ values for NVS-PAK1-1, indicating that the mutation does not alter tracer or inhibitor binding (Figure 3A, B). Therefore, we developed a full-length construct containing the Y429C mutation for NanoBRET assay development. Indeed, due to reduced binding from the AID caused by the mutation, the full-length form of the kinase demonstrated a significant BRET signal that met the threshold for assay development (Figure 3A, B). The assay window produced using the full-length mutated form of PAK1 was slightly reduced compared to the truncated forms of PAK1, likely due to residual competition from the AID. The EC_50_ determination and tracer titration, however, revealed a similar PAK1 tracer EC_50_ (Figure 3A). This suite of assays allows for unprecedented insights into the cellular target engagement of WT and mutant PAK1 by inhibitors.

### Benchmarking Literature PAK1 Inhibitors

Over the past few decades, numerous medicinal chemistry campaigns have generated compounds targeting PAK1 for various disease indications^7^. Several ATP-competitive inhibitors from these programs have demonstrated impressive potency for PAK1. Many of these ATP-competitive inhibitors, however, exhibit modest selectivity in broad kinome-wide profiling. G5555, an optimized analog of FRAX1036, is very potent in PAK1 enzymatic and substrate phosphorylation assays^29^. AZ13705339, a lead compound developed by AstraZeneca, achieved similar potency to G5555 in PAK1 biochemical and cellular phosphorylation assays^30^. While G5555 and AZ13705339 are among the most selective ATP-competitive inhibitors available for the group 1 PAKs, they still inhibit several off-target kinases. Recently, MRIA-9, a more selective derivative of G5555 optimized for salt-inducible kinase (SIK) inhibition, was shown to bind and stabilize the group 1 PAKs in thermal shift assays^31^. This compound exhibits excellent kinome-wide selectivity, potently inhibiting only the SIKs and binding the group 1 PAKs based on available data. We aimed to benchmark these inhibitors in our novel NanoBRET assay to quantify their PAK1 affinity in cells and compare results with those obtained using other PAK1 assay formats.

To quantify the potency of literature inhibitors, we determined the IC_50_ and K_i,app_ value for each of our PAK1 constructs. In the WT truncated PAK1 NanoBRET assay, the K_i,app_ values were 9, 40, 140, and 30 nM for NVS-PAK1-1, G5555, MRIA-9, and AZ13705339, respectively (Figure 4A and Table 1). These values are consistent with IC_50_ values of 5, 69, and 59 nM for NVS-PAK1-1, G5555, and AZ13705339, respectively, obtained from cellular substrate phosphorylation assays previously performed^17,29,30^. Importantly, similar values were observed for most inhibitors in the NanoBRET assay using the truncated PAK1(Y429C) mutant. AZ13705339 had an increased K_i,app_ when compared with truncated WT PAK1, suggesting that the Y429C mutation influences the active site structure (Figure 4B). Y429 is located at the P+1 pocket and, thus, is a major determinant of the structure and substrate specificity of the active conformation of the protein. Intriguingly, a divergent profile of affinity values emerged for ATP-competitive compounds with the full-length PAK1(Y429C). Specifically, all orthosteric inhibitors showed a statistically significant increase in K_i,app_ values in the full-length PAK1 Y429C assay when compared with the truncated forms of PAK1 (Figure 4B).

**Table 1:**
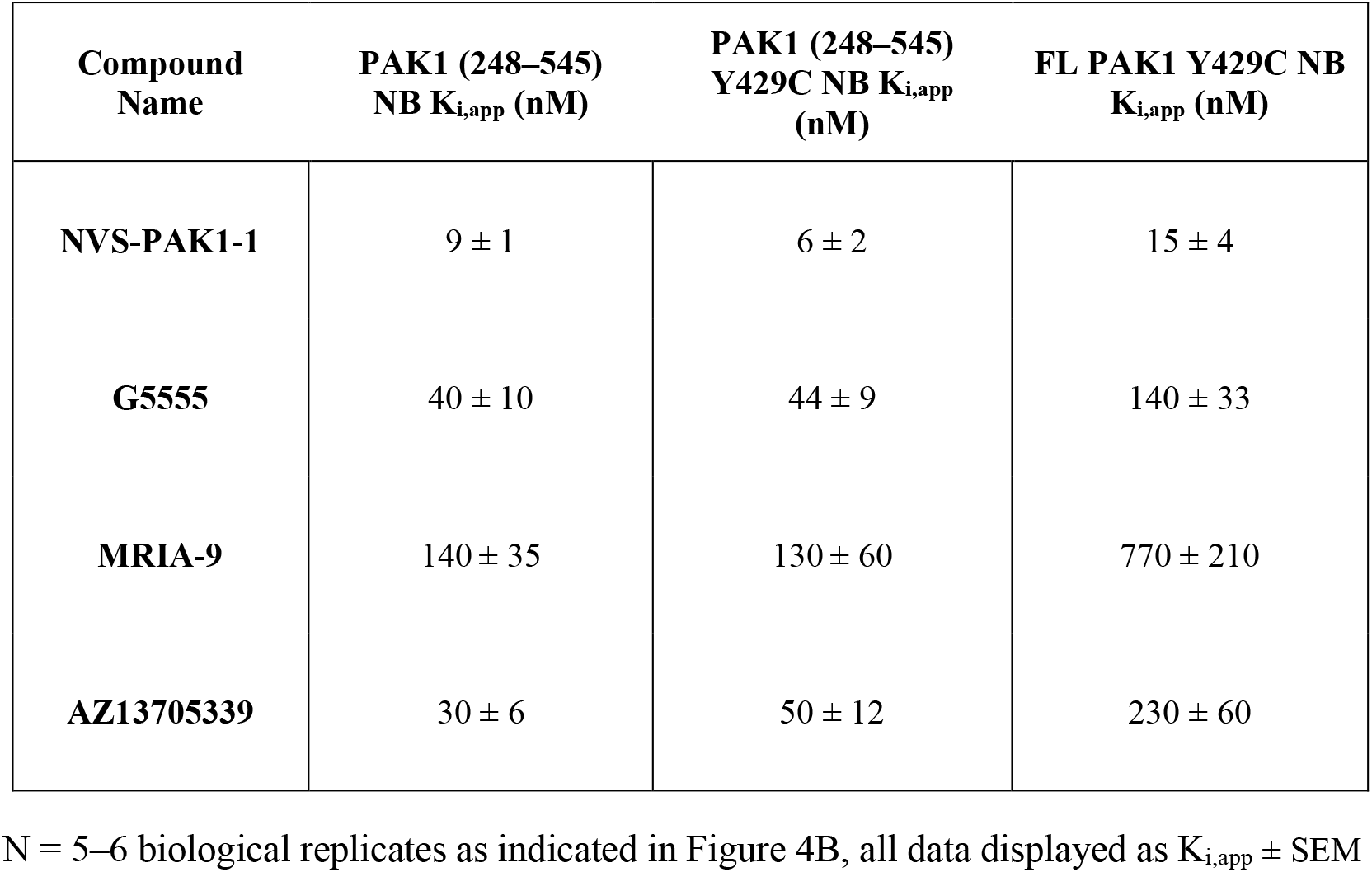
NanoBRET K_i,app_ values for literature compounds tested against various truncated and mutated forms of PAK1 in HEK293 cells.

**Figure 4.**
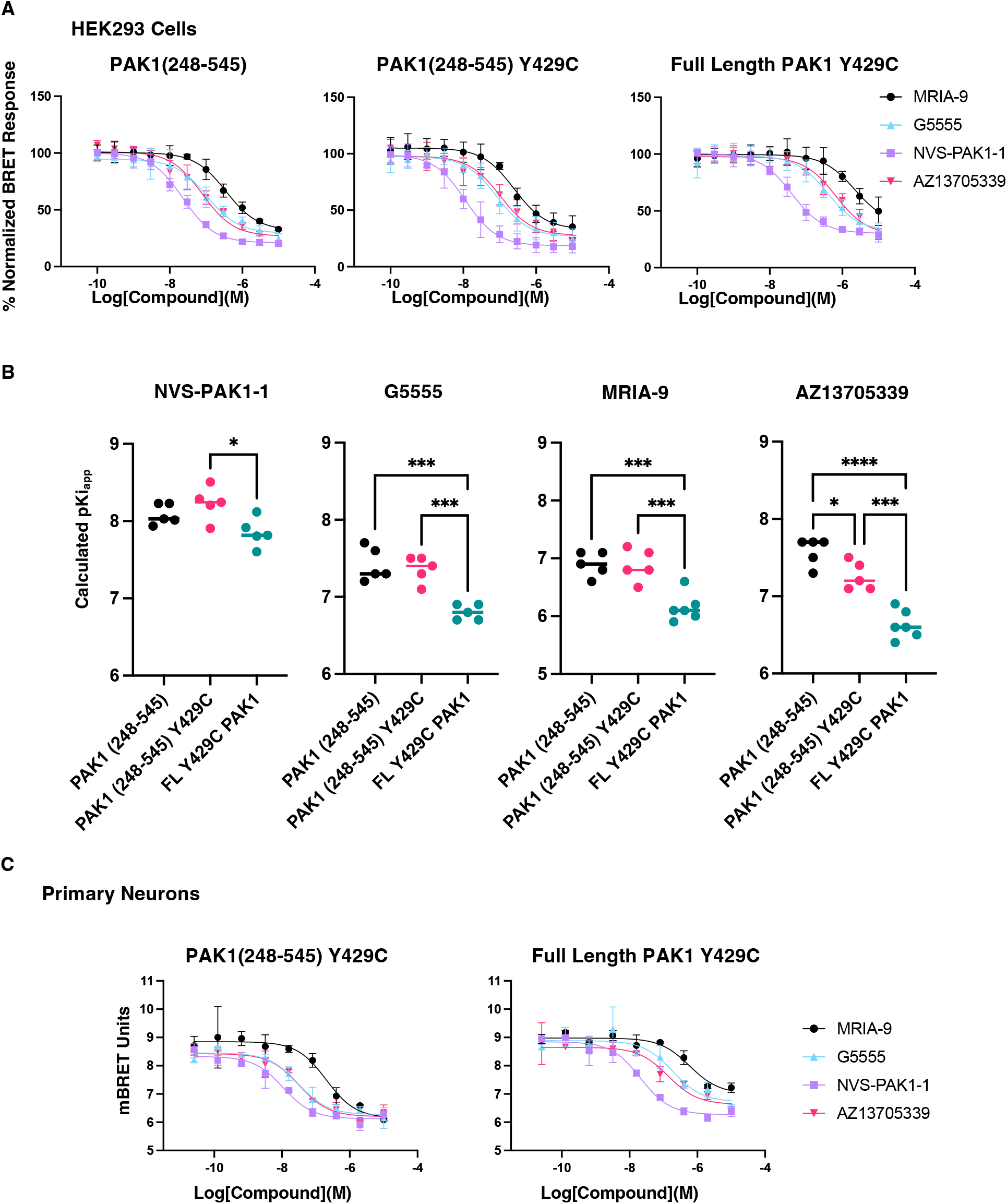
Screening of literature inhibitors in HEK293 cells and primary hippocampal neurons. (a) Normalized NanoBRET curves for literature inhibitors MRIA-9, G5555, NVS-PAK1-1, and AZ13705339.(b) Calculated cellular K_i,app_ comparison between truncated WT PAK1, truncated Y429C PAK1, and full-length Y429C PAK1. The statistical significance was assessed using a one-way ANOVA and Tukey’s post hoc analysis. For all quantifications of significance: ^****^ p < 0.0001,^***^ p < 0.001,^**^ p < 0.01, and ^*^ p < 0.05. (c) NanoBRET curves for literature inhibitors in primary neurons (n=2 ± SEM).

The altered potency observed in the presence of the N-terminal regulatory domain, which is the major structural difference between truncated and full-length PAK1(Y429C), may result from a range of effects beyond the scope of this study. The Y429C mutant does not completely ablate AID binding to the kinase domain, and approximately 30% of PAK1 proteins remain autoinhibited. Furthermore, the structural changes that underlie this reduced autoinhibition have not been explored. We speculate that, for compounds that bind the orthosteric site, the AID may sterically interfere with binding to the active site, resulting in a reduced, but not ablated, binding to the full-length form of PAK1. Consistent with this hypothesis, the shift in K_i,app_ was far more pronounced for the orthosteric inhibitors than for NVS-PAK1-1 (Figure 4B). This is harmonious with the binding mode of NVS-PAK1-1, which engages a pocket adjacent to the active site. Currently, structures of full-length PAK1 do not exist due to the disordered N-terminus of the protein. Further structural work will be required to determine the global structural changes induced by the Y429C mutation. However, the results from this assay using full-length PAK1 may better predict the true affinity for the mutated form of PAK1 that causes a neurodevelopmental disorder. Our finding is consistent with previous work showing that screening the full-length protein is vital for obtaining cellular target engagement information for a protein of interest.

### Cellular Target Engagement of PAK1 Inhibitors in Primary Neurons

To fully understand inhibitor binding, it is crucial to consider the influence of the cellular context. To date, most screening for target engagement has been performed in HEK293 cells, which may not faithfully recapitulate the environment of a target disease^32,33,34^. Indeed, differences between cell models can result in divergent target engagement profiles for the same compound^32^. To execute the optimal experiment for neurological diseases driven by PAK1 mutations, we have developed the first method to assess cellular target engagement in primary neurons. PAK1 has been implicated in several neurological maladies, including AD, Fragile X syndrome, and neurodevelopmental disorders. Using this newly developed assay, we screened NVS-PAK1-1, G5555, AZ13705339, and MRIA-9 and generated cellular K_i,app_ values similar to those observed in HEK293 cells for all PAK1 forms (Figure 4C and Table 2). Our results support that these inhibitors retain potency for mutant PAK1 in a neuronal environment and are thus viable candidates for studying neurological diseases and disorders driven by PAK1. They all support that human and rodent cells alike can be used to execute NanoBRET cellular target engagement studies.

**Table 2:** NanoBRET K_i,app_ values for literature compounds tested against various truncated and mutated forms of PAK1 in primary neurons.

| Compound Name | PAK1(248-545) Y429C NB $K_{i,app}$ (nM) | FL PAK1 Y429C NB $K_{i,app}$ (nM) |
| --- | --- | --- |
| NVS-PAK1-1 | $6 \pm 1$ | $7 \pm 1$ |
| G5555 | $16 \pm 2$ | $70 \pm 22$ |
| MRIA-9 | $100 \pm 30$ | $200 \pm 70$ |
| AZ13705339 | $20 \pm 1$ | $50 \pm 60$ |

N = 2 biological replicates, all data displayed as $K_{i,app} \pm$ SEM

## Conclusions

By synthesizing an NVS-PAK1-1-based BRET probe, we have enabled the first cellular target engagement assays for PAK1. Furthermore, we have expanded the strategy of imparting gain-of-function mutations to overcome the conformational rigidity of overexpressed proteins with autoinhibitory domains. Our assays allow for the assessment of binding to WT PAK1 using a truncated construct. The results generated with our truncated WT PAK1 construct correlate with substrate phosphorylation assays. Additionally, compounds can also be screened using either a truncated PAK1(Y429C) or a physiologically relevant full-length form of PAK1(Y429C) that drives a rare neurodevelopmental disorder. Moreover, the distinct pattern of affinity shifts observed between orthosteric inhibitors and NVS-PAK1-1 when using the full-length construct illustrates that this assay can distinguish the binding mode of PAK1 inhibitors. Finally, we have demonstrated target engagement of literature inhibitors in the first-of-its-kind PAK1 NanoBRET assay in primary hippocampal neurons. This work expands the available assays to study PAK1-driven diseases and confirms NVS-PAK1-1 as an ideal candidate for their treatment.

## Materials and Methods

### Experimental Section

#### Chemistry: General Information

All reagents and solvents that were purchased were used without further characterization or purification. Reactions listed without a reported temperature were conducted at room temperature (25°C). All reaction temperatures are listed in degrees Celsius (°C). Removal of solvent was achieved using a rotary evaporator with reduced pressure. In the reaction schemes and experimental procedures, the following abbreviations are used: min (minutes), hrs (hours), equiv (equivalents(s)), r.t. (room temperature), mg (milligrams), nmol (nanomoles), μmol (micromoles), and mmol (millimoles). The determination of compound purity and identity was achieved using microanalytical data and ^1^H NMR for intermediates and final compounds. All ^1^H NMR and ^13^C NMR spectra were recorded in a Bruker instrument using either DMSO-*d*_6_ or CD_3_OD. The magnet strength used for the NMR spectra is reported in each line listing. The peak positions are reported in parts per million (ppm) and are calibrated to the indicated deuterated solvent. The coupling constants (*J* values) are reported as the following: singlet (s), doublet (d), doublet of doublets/triplets/quartets (dd/dt/dq), doublet of doublet of doublets/triplets (ddd/ddt), triplet of doublet of triplets (tdt), triplet (t), triplet of doublets/triplets (td/tt), quartet (q), quartet of doublets (qd), pentet (p), and multiplet (m). For preparative HPLC, an Agilent 1260 Infinity II LC System equipped with a Phenomenex C18 Phenyl-Hexyl column (30 °C, 5 μm particle size, 75 × 30 mm) or an Agilent 1100 Series System equipped with a Phenomenex Luna Phenyl-Hexyl column (5 μm particle size, 100 Å pore size, 75 × 30 mm) was used. LCMS Analysis was performed using an Agilent 1290 Infinity II LC System equipped with an Agilent Infinity Lab PoroShell 120 EC-C18 column (30°C, 2.7 μm particle size, 2.1 × 50 mm), eluent 10™90% ACN in water with 0.2% formic acid or acetic acid (v/v) as indicated, and a flow rate of 1 mL/min was used. HPLC analysis confirmed that all reported compounds are >95% pure.

#### General Procedure A

To a solution of (*S*)-(3-((2-chloro-5-(2,2-difluoroethyl)-8-fluoro-5*H*-dibenzo[*b,e*][1,4]diazepin-11-yl)amino)pyrrolidin-1-yl)(1*H*-imidazol-1-yl)methanone (1.0 equiv.) in ACN, iodomethane (4.0 equiv.) was added and stirred overnight at r.t. The resulting reaction mixture was then concentrated to provide a yellow oil. The crude product was then resuspended in DCM with TEA (2.5 equiv.) and an amine-containing linker (1.0 equiv.). The reaction mixture was stirred overnight at r.t. The crude material was then concentrated *in vacuo* and purified by column chromatography on silica gel (0–10% methanol/dichloromethane) or by reverse-phase column chromatography (0–100% methanol/water) to afford the final intermediates (3–44% yield over two steps).

#### General Procedure B

The *N*-Boc-protected intermediate (excess) was resuspended in DCM (1 mL) and TFA (20 equiv.). The reaction was stirred at r.t. for two hours and then dried *in vacuo*. An excess of the crude deprotected material was then added to a vial containing DMF, DIPEA (6.0 equiv.), and 2,5-dioxopyrrolidin-1-yl 3-(5,5-difluoro-7-(1*H*-pyrrol-2-yl)-5*H*-5λ^4^,6λ^4^-dipyrrolo[1,2-*c*:2’,1’-*f*][1,3,2]diazaborinin-3-yl)propanoate (1.0 equiv.). The solution was then stirred o.n., concentrated *in vacuo*, and purified using preparative HPLC (10-100% methanol in H_2_0 + 0.05% TFA) to afford the double TFA form of the pure final product (30– 80% yield over two steps).

#### (*S*)-(3-((2-chloro-5-(2,2-difluoroethyl)-8-fluoro-5*H*-dibenzo[*b,e*][1,4]diazepin-11-yl)amino)pyrrolidin-1-yl)(1H-imidazol-1-yl)methanone (2)

This reaction was prepared according to literature precedent^21^. The reaction was carried out in a stirred solution of DCM (5 mL) containing (*S*)-2-chloro-5-(2,2-difluoroethyl)-8-fluoro-N-(pyrrolidin-3-yl)-5*H*-dibenzo[*b,e*][1,4]diazepin-11-amine (1) (300 mg, 0.76 mmol, 1.0 equiv) and carbonyldiimidazole (136 mg, 0.836 mmol, 1.1 equiv) at 0°C. To this solution, triethylamine (106 uL, 76.9 mg, 0.76 mmol, 1.0 equiv) was added, and the reaction was then stirred at r.t. overnight. The reaction was then diluted with 5 mL of water, and the organic layer was extracted with 10 mL of DCM. The organic layer was washed with brine and concentrated *in vacuo*. The crude material was purified by silica gel chromatography (0–10% methanol/dichloromethane) to afford the intermediate as a clear wax (285.7 mg, 77% yield). ^1^H NMR (400 MHz, CD_3_OD) δ 8.21 (d, *J* = 15.7 Hz, 1H), 7.59 (d, *J* = 17.3 Hz, 1H), 7.54 – 7.43 (m, 2H), 7.25 (dd, *J* = 9.0, 4.9 Hz, 1H), 7.10 (td, *J* = 10.4, 5.4 Hz, 2H), 6.74 (tt, *J* = 8.6, 2.8 Hz, 2H), 5.88 (tt, *J* = 55.9, 4.2 Hz, 1H), 4.78 – 4.63 (m, 1H), 4.20 – 3.87 (m, 4H), 3.86 – 3.70 (m, 2H), 2.49 – 2.18 (m, 2H).^1^H NMR Purity >95%. LCMS calcd for C_23_H_20_ClF_3_N_6_O [M + H]^+^: 489.1; found, 489.4.

#### *tert*-butyl (*S*)-(8-(3-((2-chloro-5-(2,2-difluoroethyl)-8-fluoro-5*H*-dibenzo[*b,e*][1,4]diazepin-11-yl)amino)pyrrolidine-1-carboxamido)octyl)carbamate (3)

The reaction was performed according to general procedure A with (*S*)-(3-((2-chloro-5-(2,2-difluoroethyl)-8-fluoro-5*H*-dibenzo[*b,e*][1,4]diazepin-11-yl)amino)pyrrolidin-1-yl)(1*H*-imidazol-1-yl)methanone (29 mg, 59 μmol) in ACN (1 mL) with iodomethane (34 mg, 240 μmol). The crude material was then reacted with DCM (2 mL) and *tert*-butyl (8-aminooctyl)carbamate (15 mg, 60 μmol) in the presence of triethylamine (15 mg, 150 μmol). The concentrated reaction mixture was purified by reverse-phase column chromatography (0–100% methanol/water) to afford the acetic acid salt intermediate as a colorless powder (17.6 mg, 44% yield over two steps). ^1^H NMR (500 MHz, cd_3_od) δ 7.48 – 7.36 (m, 2H), 7.24 (dd, *J* = 8.7, 2.2 Hz, 1H), 7.08 (dt, *J* = 8.9, 5.3 Hz, 1H), 6.82 – 6.66 (m, 2H), 5.87 (tt, *J* = 55.7, 4.2 Hz, 1H), 4.65 (dp, *J* = 27.0, 5.6 Hz, 1H), 4.16 – 3.95 (m, 2H), 3.78 (dt, *J* = 10.5, 6.4 Hz, 1H), 3.57 (ddt, *J* = 14.9, 10.4, 5.9 Hz, 2H), 3.51 – 3.42 (m, 1H), 3.40 – 3.34 (m, 1H), 3.26 – 3.12 (m, 2H), 3.01 (q, *J* = 6.8 Hz, 2H), 2.31 (ddq, *J* = 33.3, 13.4, 6.9 Hz, 1H), 2.23 – 2.03 (m, 1H), 1.53 (h, *J* = 7.5 Hz, 3H), 1.45 (s, 12H), 1.34 (s, 6H), 1.30 (d, *J* = 5.3 Hz, 2H). ^1^H NMR Purity >95%. LCMS calcd for C_33_H_44_ClF_3_N_6_O_3_ [M + H]^+^: 665.3; found, 665.3.

#### *tert*-butyl (*S*)-(12-(3-((2-chloro-5-(2,2-difluoroethyl)-8-fluoro-5*H*-dibenzo[*b,e*][1,4]diazepin-11-yl)amino)pyrrolidine-1-carboxamido)dodecyl)carbamate (4)

The reaction was performed according to general procedure A with (*S*)-(3-((2-chloro-5-(2,2-difluoroethyl)-8-fluoro-5*H*-dibenzo[*b,e*][1,4]diazepin-11-yl)amino)pyrrolidin-1-yl)(1*H*-imidazol-1-yl)methanone (29 mg, 59 μmol) in ACN (1 mL) with iodomethane (34 mg, 240 μmol). The crude material was then reacted with DCM (2 mL) and *tert*-butyl (12-aminododecyl)carbamate (119 mg, 397 μmol) in the presence of DIPEA (100 mg, 992 μmol). The concentrated reaction mixture was purified by reverse-phase column chromatography (0–100% methanol/water) to afford the acetic acid salt intermediate as a colorless powder (9 mg, 3% yield over two steps). ^1^H NMR (500 MHz, cd_3_od) δ 7.48 – 7.36 (m, 2H), 7.24 (dd, *J* = 8.7, 1.6 Hz, 1H), 7.12 – 7.05 (m, 1H), 6.84 – 6.66 (m, 2H), 6.05 – 5.69 (m, 1H), 4.65 (dp, *J* = 26.8, 5.7 Hz, 1H), 4.16 – 3.95 (m, 2H), 3.77 (dt, *J* = 11.7, 5.9 Hz, 1H), 3.63 – 3.52 (m, 2H), 3.51 – 3.43 (m, 1H), 3.37 (dd, *J* = 10.5, 4.7 Hz, 1H), 3.26 – 3.12 (m, 2H), 3.03 (t, *J* = 7.6 Hz, 2H), 2.31 (ddq, *J* = 33.6, 13.6, 7.0 Hz, 1H), 2.22 – 2.06 (m, 1H), 1.52 (p, *J* = 7.6 Hz, 3H), 1.45 (s, 11H), 1.37 – 1.30 (m, 11H), 1.27 (s, 5H).^1^H NMR Purity >95%. LCMS calcd for C_37_H_52_ClF_3_N_6_O_3_ [M + H]^+^: 721.4; found, 721.8.

#### *tert*-butyl (*S*)-(1-(3-((2-chloro-5-(2,2-difluoroethyl)-8-fluoro-5*H*-dibenzo[*b,e*][1,4]diazepin-11-yl)amino)pyrrolidin-1-yl)-1-oxo-5,8,11-trioxa-2-azatridecan-13-yl)carbamate (5)

The reaction was performed according to general procedure A with (*S*)-(3-((2-chloro-5-(2,2-difluoroethyl)-8-fluoro-5*H*-dibenzo[*b,e*][1,4]diazepin-11-yl)amino)pyrrolidin-1-yl)(1*H*-imidazol-1-yl)methanone (194 mg, 397 μmol) in ACN (2 mL) with iodomethane (225 mg, 1.59 mmol). The crude material was then reacted with DCM (2 mL) and *tert*-butyl (2-(2-(2-(2-aminoethoxy)ethoxy)ethoxy)ethyl)carbamate (116 mg, 397 μmol) in the presence of triethylamine (100 mg, 992 μmol). The concentrated reaction mixture was purified by silica gel chromatography (0–10% methanol/dichloromethane) to afford the intermediate as a clear wax (15 mg, 5.3% yield over two steps). ^1^H NMR (500 MHz, cd_3_od) δ 7.76 – 7.66 (m, 2H), 7.53 (d, *J* = 8.8 Hz, 1H), 7.49 (dd, *J* = 9.0, 5.2 Hz, 1H), 7.24 – 7.15 (m, 2H), 6.02 (tq, *J* = 55.2, 3.5 Hz, 1H), 4.71 (ddt, *J* = 7.4, 5.6, 2.8 Hz, 1H), 4.41 – 4.18 (m, 2H), 3.92 – 3.46 (m, 19H), 3.39 (dt, *J* = 9.8, 5.6 Hz, 2H), 3.23 (td, *J* = 5.7, 2.8 Hz, 2H), 2.25 (ddt, *J* = 12.1, 7.8, 4.2 Hz, 2H), 1.44 (d, *J* = 3.1 Hz, 9H). ^1^H NMR Purity >95%. LCMS calcd for C_33_H_44_ClF_3_N_6_O_6_ [M + H]^+^: 713.3; found, 713.3.

#### *tert*-butyl (*S*)-(2-(3-((2-chloro-5-(2,2-difluoroethyl)-8-fluoro-5*H*-dibenzo[*b,e*][1,4]diazepin-11-yl)amino)pyrrolidine-1-carboxamido)ethyl)carbamate (6)

The reaction was performed according to general procedure A with (*S*)-(3-((2-chloro-5-(2,2-difluoroethyl)-8-fluoro-5*H*-dibenzo[*b,e*][1,4]diazepin-11-yl)amino)pyrrolidin-1-yl)(1*H*-imidazol-1-yl)methanone (194 mg, 397 μmol) in ACN (2 mL) with iodomethane (225 mg, 1.59 mmol). The crude material was then reacted with *tert*-butyl (2-aminoethyl)carbamate (63.6 mg, 397 μmol) and triethylamine (100 mg, 992 μmol) in DCM (2 mL). The concentrated reaction mixture was purified by silica gel column chromatography (0– 10% methanol/dichloromethane) to afford the intermediate as a clear wax (43.9 mg, 19% yield over two steps). ^1^H NMR (400 MHz, cd_3_od) δ 7.71 (ddd, *J* = 8.7, 2.5, 1.7 Hz, 1H), 7.65 (d, *J* = 2.5 Hz, 1H), 7.54 – 7.42 (m, 2H), 7.21 – 7.12 (m, 2H), 5.99 (td, *J* = 55.3, 3.4 Hz, 1H), 4.68 – 4.64 (m, 1H), 4.37 – 4.19 (m, 2H), 3.90 – 3.79 (m, 1H), 3.74 – 3.41 (m, 4H), 3.20 (dt, *J* = 28.4, 6.5 Hz, 4H), 2.57 – 2.17 (m, 2H), 1.43 (s, 9H). ^1^H NMR Purity >95%. LCMS calcd for C_27_H_32_ClF_3_N_6_O_3_ [M + H]^+^: 581.2; found, 581.3.

#### (*S*)-3-((2-chloro-5-(2,2-difluoroethyl)-8-fluoro-5*H*-dibenzo[*b,e*][1,4]diazepin-11-yl)amino)-*N*-(9-(3-(5,5-difluoro-7-(1*H*-pyrrol-2-yl)-5*H*-5λ^4^,6λ^4^-dipyrrolo[1,2-*c*:2’,1’-*f*][1,3,2]diazaborinin-3-yl)propanamido)nonyl)pyrrolidine-1-carboxamide (7)

The reaction was performed according to general procedure B with *tert*-butyl (*S*)-(8-(3-((2-chloro-5-(2,2-difluoroethyl)-8-fluoro-5*H*-dibenzo[b,e][1,4]diazepin-11-yl)amino)pyrrolidine-1-carboxamido)octyl)carbamate (17.6 mg, 26.5 μmol), DCM (1 mL), and TFA (60.3 mg, 529 μmol). The crude material was reacted with DMF (1 mL), DIPEA (10 mg, 100 μmol), and 2,5-dioxopyrrolidin-1-yl 3-(5,5-difluoro-7-(1*H*-pyrrol-2-yl)-5*H*-5λ^4^,6λ^4^-dipyrrolo[1,2-*c*:2’,1’-*f*][1,3,2]diazaborinin-3-yl)propanoate (7 mg, 20 μmol). The title compound was purified to afford a purple amorphous solid (14 mg, 80% yield over two steps). ^1^H NMR (850 MHz, cd_3_od) δ 7.68 (d, *J* = 8.8 Hz, 1H), 7.61 (dd, *J* = 9.3, 2.4 Hz, 1H), 7.48 (dd, *J* = 9.2, 2.9 Hz, 1H), 7.43 (s, 1H), 7.26 (d, *J* = 1.4 Hz, 1H), 7.24 – 7.19 (m, 3H), 7.16 – 7.11 (m, 2H), 7.04 (dd, *J* = 4.5, 1.9 Hz, 1H), 6.94 (dd, *J* = 4.0, 2.0 Hz, 1H), 6.37 – 6.32 (m, 2H), 5.98 (tq, *J* = 55.3, 3.4 Hz, 1H), 4.67 – 4.62 (m, 1H), 4.30 (t, *J* = 13.4 Hz, 1H), 4.27 – 4.16 (m, 1H), 3.84 (ddd, *J* = 18.9, 11.3, 6.0 Hz, 1H), 3.71 – 3.60 (m, 1H), 3.60 – 3.47 (m, 2H), 3.29 (td, *J* = 7.5, 4.0 Hz, 2H), 3.21 – 3.14 (m, 4H), 2.66 – 2.61 (m, 2H), 2.53 – 2.40 (m, 1H), 2.39 – 2.17 (m, 1H), 1.51 (dp, *J* = 26.5, 6.7 Hz, 4H), 1.36 – 1.29 (m, 8H). HPLC Purity >95%. LCMS calcd for C_45_H_50_BClF_5_N_9_O_2_ [M + H]^+^: 876.4; found, 876.2.

#### (*S*)-3-((2-chloro-5-(2,2-difluoroethyl)-8-fluoro-5*H*-dibenzo[*b,e*][1,4]diazepin-11-yl)amino)-*N*-(12-(3-(5,5-difluoro-7-(1*H*-pyrrol-2-yl)-5*H*-5λ^4^,6λ^4^-dipyrrolo[1,2-*c*:2’,1’-*f*][1,3,2]diazaborinin-3-yl)propanamido)dodecyl)pyrrolidine-1-carboxamide (8)

The reaction was performed according to general procedure B with *tert*-butyl (*S*)-(12-(3-((2-chloro-5-(2,2-difluoroethyl)-8-fluoro-5*H*-dibenzo[*b,e*][1,4]diazepin-11-yl)amino)pyrrolidine-1-carboxamido)dodecyl)carbamate (9 mg, 20 μmol), DCM (1 mL), and TFA (30 mg, 200 μmol). The crude material was reacted with DMF (1 mL), DIPEA (9 mg, 70 μmol), and 2,5-dioxopyrrolidin-1-yl 3-(5,5-difluoro-7-(1*H*-pyrrol-2-yl)-5*H*-5λ^4^,6λ^4^-dipyrrolo[1,2-*c*:2’,1’-*f*][1,3,2]diazaborinin-3-yl)propanoate (5 mg, 10 μmol). The title compound was purified to afford a purple amorphous solid as a double TFA salt (4.5 mg, 30% yield over two steps). ^1^H NMR (400 MHz, cd_3_od) δ 7.66 (dd, *J* = 8.8, 2.5 Hz, 1H), 7.59 (dd, *J* = 5.6, 2.5 Hz, 1H), 7.45 (d, *J* = 8.8 Hz, 1H), 7.42 – 7.37 (m, 1H), 7.29 – 7.15 (m, 4H), 7.14 – 7.05 (m, 2H), 7.01 (d, *J* = 4.6 Hz, 1H), 6.91 (d, *J* = 4.0 Hz, 1H), 6.37 – 6.29 (m, 2H), 5.95 (t, *J* = 55.3 Hz, 1H), 4.62 (d, *J* = 4.0 Hz, 1H), 4.40 – 4.12 (m, 2H), 3.82 (td, *J* = 10.2, 6.0 Hz, 1H), 3.71 – 3.44 (m, 3H), 3.27 (d, *J* = 7.7 Hz, 2H), 3.21 – 3.09 (m, 4H), 2.62 (t, *J* = 7.7 Hz, 2H), 1.57 – 1.43 (m, 4H), 1.39 – 1.20 (m, 18H). HPLC Purity >95%. LCMS calcd for C_48_H_56_BClF_5_N_9_O_2_ [M + H]^+^: 932.4; found, 932.9.

#### (*S*)-3-((2-chloro-5-(2,2-difluoroethyl)-8-fluoro-5*H*-dibenzo[*b,e*][1,4]diazepin-11-yl)amino)-*N*-(15-5,5-difluoro-7-(1*H*-pyrrol-2-yl)-5*H*-5λ^4^,6λ^4^-dipyrrolo[1,2-*c*:2’,1’-*f*][1,3,2]diazaborinin-3-yl)-13-oxo-3,6,9-trioxa-12-azapentadecyl)pyrrolidine-1-carboxamide (9)

The reaction was performed according to general procedure B with *tert*-butyl (*S*)-(1-(3-((2-chloro-5-(2,2-difluoroethyl)-8-fluoro-5*H*-dibenzo[*b,e*][1,4]diazepin-11-yl)amino)pyrrolidin-1-yl)-1-oxo-5,8,11-trioxa-2-azatridecan-13-yl)carbamate (15 mg, 21 μmol), DCM (1 mL), and TFA (48 mg, 420 μmol). The crude material was reacted with DMF (1 mL), DIPEA (10 mg, 100 μmol), and 2,5-dioxopyrrolidin-1-yl 3-(5,5-difluoro-7-(1*H*-pyrrol-2-yl)-5*H*-5λ^4^,6λ^4^-dipyrrolo[1,2-*c*:2’,1’-*f*][1,3,2]diazaborinin-3-yl)propanoate (8 mg, 20 μmol). The title compound was purified to afford a purple amorphous solid as a double TFA salt (3.9 mg, 20% yield over two steps). ^1^H NMR (850 MHz, cd_3_od) δ 7.67 (dt, *J* = 8.8, 2.2 Hz, 1H), 7.60 (d, *J* = 2.5 Hz, 1H), 7.47 (dd, *J* = 8.9, 2.3 Hz, 1H), 7.43 (ddd, *J* = 7.5, 5.0, 3.0 Hz, 1H), 7.24 (d, *J* = 1.8 Hz, 1H), 7.22– 7.17 (m, 3H), 7.14 (ddt, *J* = 10.9, 8.3, 2.6 Hz, 2H), 7.02 (dd, *J* = 4.5, 3.1 Hz, 1H), 6.93 (t, *J* = 3.7 Hz, 1H),6.36 – 6.31 (m, 2H), 5.96 (td, *J* = 3.6, 1.4 Hz, 1H), 4.64 (ddt, *J* = 10.3, 5.7, 3.7 Hz, 1H), 4.33 – 4.25 (m,1H), 4.20 (qdd, *J* = 14.8, 6.7, 3.4 Hz, 1H), 3.81 (ddd, *J* = 17.4, 11.4, 6.0 Hz, 1H), 3.68 (dd, *J* = 11.0, 3.3Hz, 1H), 3.64 – 3.57 (m, 9H), 3.56 – 3.43 (m, 6H), 3.40 – 3.32 (m, 4H), 3.29 – 3.22 (m, 2H), 2.68 – 2.61(m, 2H), 2.43 (ddtd, *J* = 49.6, 14.0, 8.2, 6.0 Hz, 1H), 2.36 – 2.16 (m, 1H). HPLC Purity >95%. LCMS calcd for C_44_H_48_BClF_5_N_9_O_5_ [M + H]^+^: 924.8; found, 924.8.

#### (*S*)-3-((2-chloro-5-(2,2-difluoroethyl)-8-fluoro-5*H*-dibenzo[*b,e*][1,4]diazepin-11-yl)amino)-*N*-(2-(3-(5,5-difluoro-7-(1*H*-pyrrol-2-yl)-5*H*-5λ^4^,6λ^4^-dipyrrolo[1,2-*c*:2’,1’-*f*][1,3,2]diazaborinin-3-yl)propanamido)ethyl)pyrrolidine-1-carboxamide (10)

The reaction was performed according to general procedure B with *tert*-butyl (*S*)-(2-(3-((2-chloro-5-(2,2-difluoroethyl)-8-fluoro-5*H*-dibenzo[*b,e*][1,4]diazepin-11-yl)amino)pyrrolidine-1-carboxamido)ethyl)carbamate (43.9 mg, 75.6 μmol), DCM (1 mL), and TFA (172 mg, 1.51 mmol). The crude material was reacted with DMF (1 mL), DIPEA (10 mg, 80 μmol), and 2,5-dioxopyrrolidin-1-yl 3-(5,5-difluoro-7-(1*H*-pyrrol-2-yl)-5*H*-5λ^4^,6λ^4^-dipyrrolo[1,2-*c*:2’,1’-*f*][1,3,2]diazaborinin-3-yl)propanoate (6 mg, 10 μmol). The title compound was purified to afford a purple amorphous solid as a double TFA salt (4.3 mg, 30% yield over two steps). ^1^H NMR (400 MHz, cd_3_od) δ 7.69 – 7.58 (m, 2H), 7.47 – 7.36 (m, 2H),7.27 – 6.98 (m, 7H), 6.92 (t, *J* = 4.4 Hz, 1H), 6.36 – 6.24 (m, 2H), 5.92 (tdt, *J* = 55.3, 9.8, 3.6 Hz, 1H), 4.62– 4.54 (m, 1H), 4.31 – 4.10 (m, 2H), 3.77 (ddd, *J* = 13.0, 11.4, 5.7 Hz, 1H), 3.71 – 3.55 (m, 1H), 3.53 –3.32 (m, 4H), 3.29 – 3.14 (m, 4H), 2.70 – 2.56 (m, 2H), 2.39 (tt, *J* = 14.2, 7.7 Hz, 1H), 2.32 – 2.13 (m, 1H). HPLC Purity >95%. LCMS calcd for C_38_H_36_BClF_5_N_9_O_2_ [M + H]^+^: 792.3; found, 792.7.

## Biology Methods

### General information for NanoBRET Assays in Human Embryonic Kidney (HEK293) cells

All NanoBRET assays carried out in HEK293 cells were performed using a modified version of existing literature protocols^21,23,25^. HEK293 cells were purchased from ATCC and cultured in Dulbecco’s modified Eagle medium (DMEM; Gibco) supplemented with 10% fetal bovine serum (FBS; Avantor) at 37 °C in 5% CO_2_. For transfections, a DNA transfection complex was formed with 10 μg/mL of DNA in Opti-MEM without serum or phenol red (Gibco) and 30 μL/mL of FuGENE HD (Promega). To create the DNA solution, 9 μg/mL of carrier DNA (Promega) and 1 μg/mL NLuc C-terminal fusion protein (PAK1(248–545)-NLuc, PAK1(248–545) (Y429C)-NLuc, or PAK1(Y429C)-NLuc). Upon addition of FuGENE HD to the DNA solution, the mixture was vortexed and incubated at room temperature (r.t.) for 20 minutes. A 20x volume of HEK293 cells suspended in DMEM with 10% FBS was added to the transfection complex solution to create a final concentration of 2×10^5^ cells/mL. 100 μL of the final cell solution was transferred to a 96-well tissue culture-treated plate (Corning).

After 24 hours of incubation at 37 °C in 5% CO_2_, the media on the transfected cells was aspirated and replaced with 90 μL of Opti-MEM in the negative control wells and 85 μL in wells with tracer added. Then, 5 μL of a 20x tracer dilution solution was added to every well, excluding the negative control wells. For control wells, 10 μL of Opti-MEM with DMSO was added. For test wells, 10 μL of a 10x compound stock solution made up in Opti-MEM was added. After all additions, each well contained 100 μL and 1.1% DMSO. After compound and tracer addition, the plate was then incubated for 2 hours at 37 °C in 5% CO_2_. For the luciferase substrate solution, the NanoBRET NanoGLO substrate (Promega) was added at a ratio of 1:166 in Opti-MEM and extracellular NLuc inhibitor (Promega) was added at a ratio of 1:500. 50 μL of the luciferase substrate solution was added to each well of the plate after they had been cooled to r.t. The plate was then read on a GloMAX Discover luminometer (Promega) equipped with a 450 nm BP filter (donor) and a 600 nm LP filter (acceptor) within 10 minutes of substrate addition. To produce milliBRET units (mBU), the acceptor emissions (600 nm) were divided by the donor emissions (450 nm) and the result was multiplied by 1000. All mBU values in this study are background corrected by subtracting the average of the negative control wells, unless otherwise indicated. NanoBRET values were normalized to % BRET response by dividing background-subtracted values of each test well by the top control (tracer only) wells and multiplying by 100.

### NanoBRET Tracer Competition Experiments

A 20x stock of tracer **10** was prepared by dilution such that the final concentrations were 500, 400, 300, or 200 nM. 5 μL of the 20x tracer solution was added to each test well in a 96-well plate. NVS-PAK1-1 was tested in an 11-point dose–response format against all the concentrations of tracer. The 3 biological replicates were plotted using a sigmoidal three-parameter dose–response logistical curve in the GraphPad Prism software. The error bars indicated the standard error of the mean.

### NanoBRET Tracer EC_50_ Determination

Tracer **10**, prepared in a dilution series with tracer dilution buffer and DMSO, was tested in an 11-point dose–response fashion in cells transfected with the indicated NLuc fusion protein. Three biological replicates were plotted using a sigmoidal three-parameter dose–response logistical curve in the GraphPad Prism software. The error bars indicated the standard error of the mean.

### NanoBRET Inhibitor Screening Assay

A 20x stock solution was prepared for each NLuc fusion protein, with the final concentration being 500 nM for PAK1(248–545) and full-length PAK1 Y429C, and 250 nM for PAK1(248–545) Y429C. Each compound was tested in an 11-point dose–response format by adding a 10 μL of a 10x dilution series prepared in Opti-MEM. The data were plotted using a Sigmoidal three-parameter dose–response logistical curve in the GraphPad Prism software.

### Primary Neuron Culture and Seeding

E18 rat hippocampal neurons were isolated and cultured according to established protocols^35^. Neurons were seeded at a density of 100,000 cells per well in 96-well plates pre-coated with laminin (10 µg/mL, Corning, Ref #354232) and poly-D-lysine (0.5 mg/mL, Sigma, Cas #27964-99-4). Neurons were maintained in NbActiv4 medium without phenol red (BrainBits, NB4PR500) and allowed to adhere at 37°C and 5% CO_2_ for 24 hours prior to transduction.

### Determination of Optimal MOI

To determine the optimal multiplicity of infection (MOI), neurons were transduced with lentiviral constructs encoding either PAK1(248–545) (Y429C)-NLuc (Vector Builder, VB240819-1550ewj) or PAK1(Y429C)-NLuc (full-length, Vector Builder, VB240819-1545mzh) (Supplementary Figure S2) with varying MOI. Following a 24-hour incubation period, the media were removed and replaced by 100 µL of fresh NbActiv4 medium without phenol red. Cells were then incubated for an additional 72 hours. The plates were then taken out of the incubator, and a complete NanoBRET substrate (Promega), containing extracellular NLuc inhibitor at 20 µM (Promega, Ref #N235B) and NanoBRET Nano-Glo Substrate at 1:166 (Promega, Ref #N157C) was added and incubated for 2–3 minutes. The optimal MOI was defined as the lowest MOI that results in ≥80% of the maximum signal.

### NanoBRET Inhibitor Screening Assay for PAK1 in Neurons

Neurons were transduced with the lentiviral constructs at an MOI of 50 in NbActiv4 medium without phenol red. After 24 hours, the viral medium was replaced with fresh medium, and cells were incubated for an additional 72 hours. Tracer **10** was added to the neurons at a final concentration of 500 nM and 250 nM for cells transduced with PAK1(Y429C)-NLuc and PAK1(248–545) (Y429C)-NLuc, respectively. The cells were immediately treated with PAK1 inhibitors at a concentration gradient from 10 to 0.0000256 µM and incubated for 2 hours at 37°C and 5% CO_2_. Following incubation, the NanoBRET NanoGLO substrate was added for 2–3 minutes. Donor emission was measured at a wavelength of 450 nm, and acceptor emission at 630 nm using the GloMAX Discover luminometer (Promega).

## Author Contributions

All authors have given approval to the final version of the manuscript.

## Notes

The authors declare no competing financial interest.

## Acknowledgment

The NanoBRET construct for PAK1(248-545)-NLuc used in this study was generously provided by Promega Corporation (Madison, WI). We thank Matthew Robers for his helpful insight on this project. We would also like to acknowledge the University of Miami TMP Drug Discovery Core (RRID: SCR_022542). Table of contents graphic created with BioRender.com.

## Abbreviation

ACN: acetonitrile
AD: Alzheimer’s disease
AID: autoinhibitory domain
ATP: adenosine 5′-triphosphate
Boc: *tert*-butyloxycarbonyl
BRET: bioluminescence resonance energy transfer
CDC42: cell division cycle 42
CDI: 1,1′-carbonyldiimidazole
cd_3_od: methanol-*d*_4_
DCM: dichloromethane
DIPEA: *N,N*-diisopropylethylamine
DMEM: Dulbecco’s Modified Eagle Medium
DMF: *N,N*-dimethylformamide
DMSO: dimethyl sulfoxide
DMSO-*d*_6_: dimethyl sulfoxide-*d*_6_
DNA: deoxyribonucleic acid
EC_50_: half-maximal effective concentration
FBS: fetal bovine serum
FL: full-length
HATU: 1-[bis(dimethylamino)methylene]-1*H*-1,2,3-triazolo[4,5-*b*]pyridinium 3-oxide hexafluorophosphate
HEK293: human embryonic kidney 293
HPLC: high-performance liquid chromatography
IC_50_: half-maximal inhibitory concentration
*K*_i_,_app_: apparent inhibition constant
*K*_m_: Michaelis constant
LCMS: liquid chromatography–mass spectrometry
MAPK: mitogen-activated protein kinase
mBU: milliBRET units
MeI: methyl iodide
MOI: multiplicity of infection
NanoBRET: nanoluciferase-based bioluminescence resonance energy transfer
NB: NanoBRET
NLuc: Nanoluciferase
NMR: nuclear magnetic resonance
PAK: p21-activated kinase
PEG: polyethylene glycol
PROTAC: proteolysis-targeting chimera
Rac1: Ras-related C3 botulinum toxin substrate 1
SEM: standard error of the mean
SGC: Structural Genomics Consortium
SIK: salt-inducible kinase
SYK: spleen tyrosine kinase
TEA: triethylamine
TFA: trifluoroacetic acid
WNT: wingless-related integration site
WT: wild-type.

