## Supplementary Information for "Development of NanoBRET cellular target engagement assays in primary neurons for activating mutants of p21-activated kinase 1"

### Table of Contents

Figure S1 S2

Figure S2 S3

Purity traces and spectra for all compounds S4–S10

**Figure S1:**

**A**

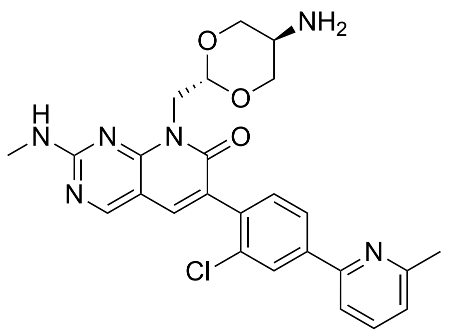

G5555
Group 1 PAK inhibitor
PAK1 $IC_{50}$
3.7 nM biochemical
69 nM Cellular (pMEK)

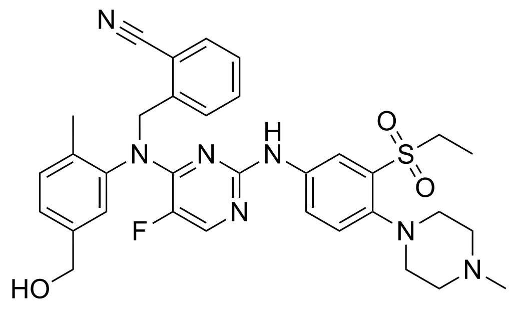

AZ13705339
Group 1 PAK inhibitor
PAK1 $IC_{50}$
0.33 nM biochemical
59 nM Cellular (pPAK1)

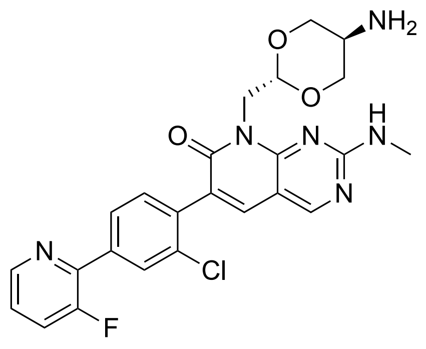

MRIA-9
Pan SIK inhibitor
PAK1 Thermal Shift
4.9°C (DSF)

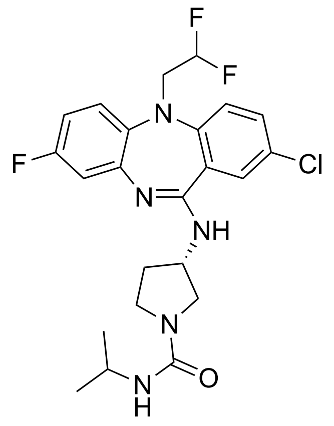

NVS-PAK1-1
PAK1 Chemical Probe
PAK1 $IC_{50}$
7 nM (biochemical)
5.2 nM (pPAK1)

**Supplementary Figure 1:** Chemical structure of all tested PAK1 inhibitors with a brief description and the associated PAK1 affinity metrics.

**Figure S2:**

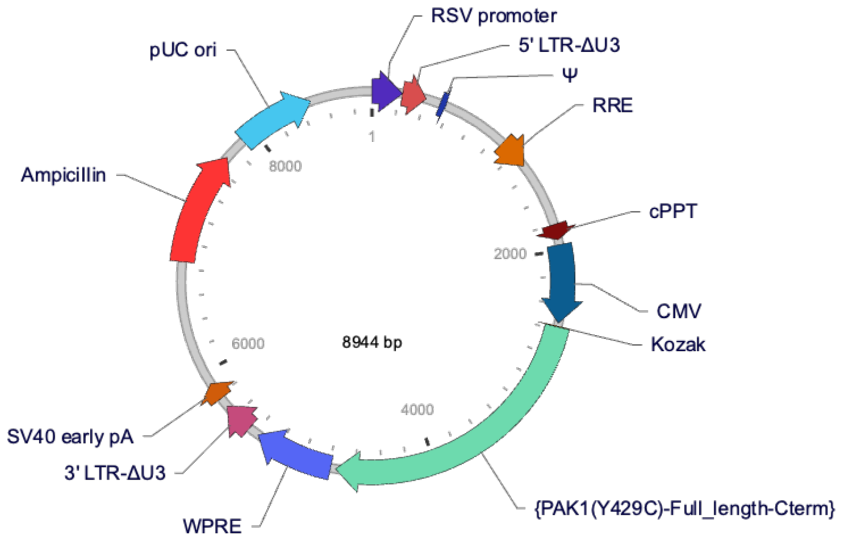


pLV[Exp]-CMV>{PAK1(Y429C)-Full-length-Cterm}

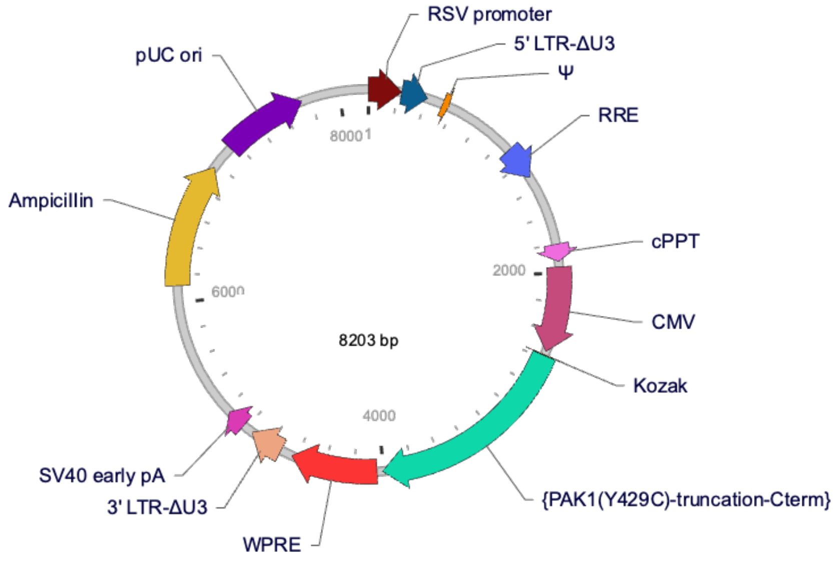


pLV[Exp]-CMV>{PAK1(Y429C)-truncation-Cterm}

**Supplementary Figure 2: Vector Structure of PAK1 lentivirus used for Neuronal transduction.**

[1]H NMR (400 MHz, $CD_3OD$) for compound **2**

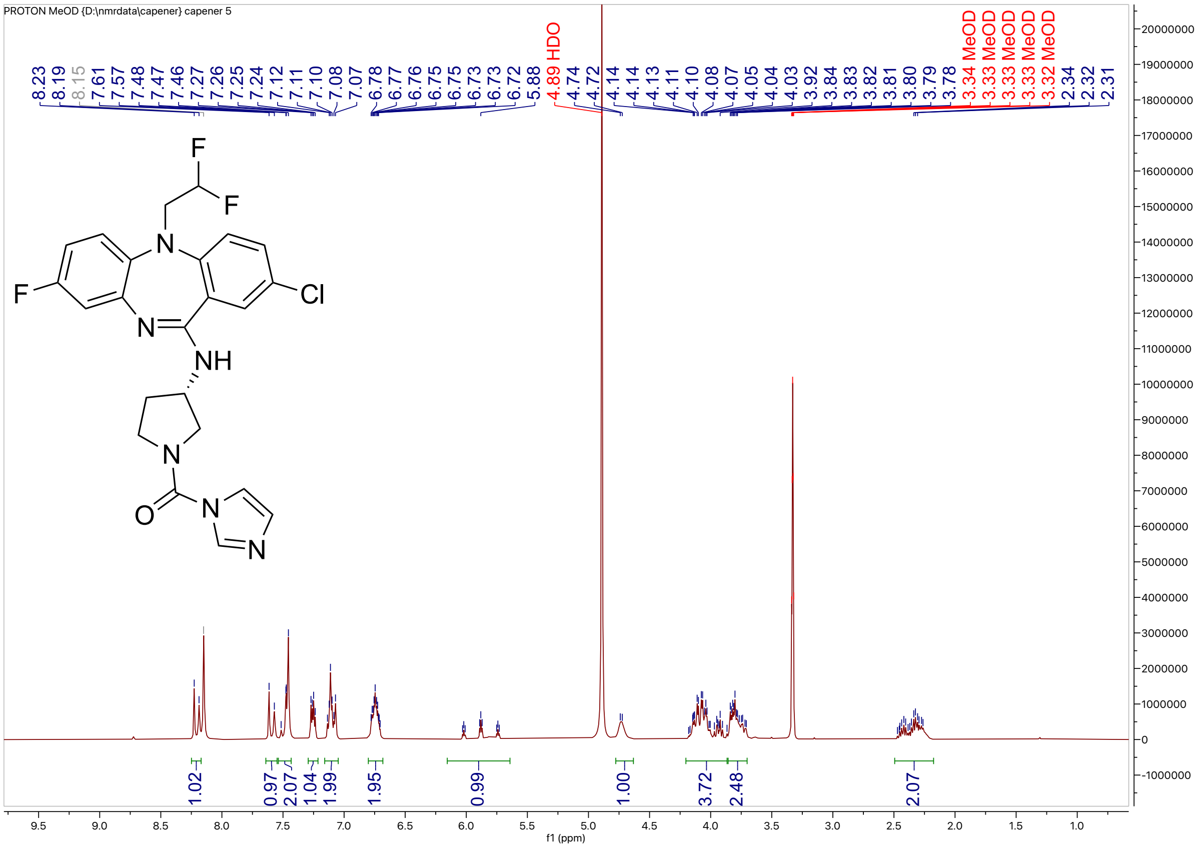


[1]H NMR (400 MHz, $CD_3OD$) for compound **3**

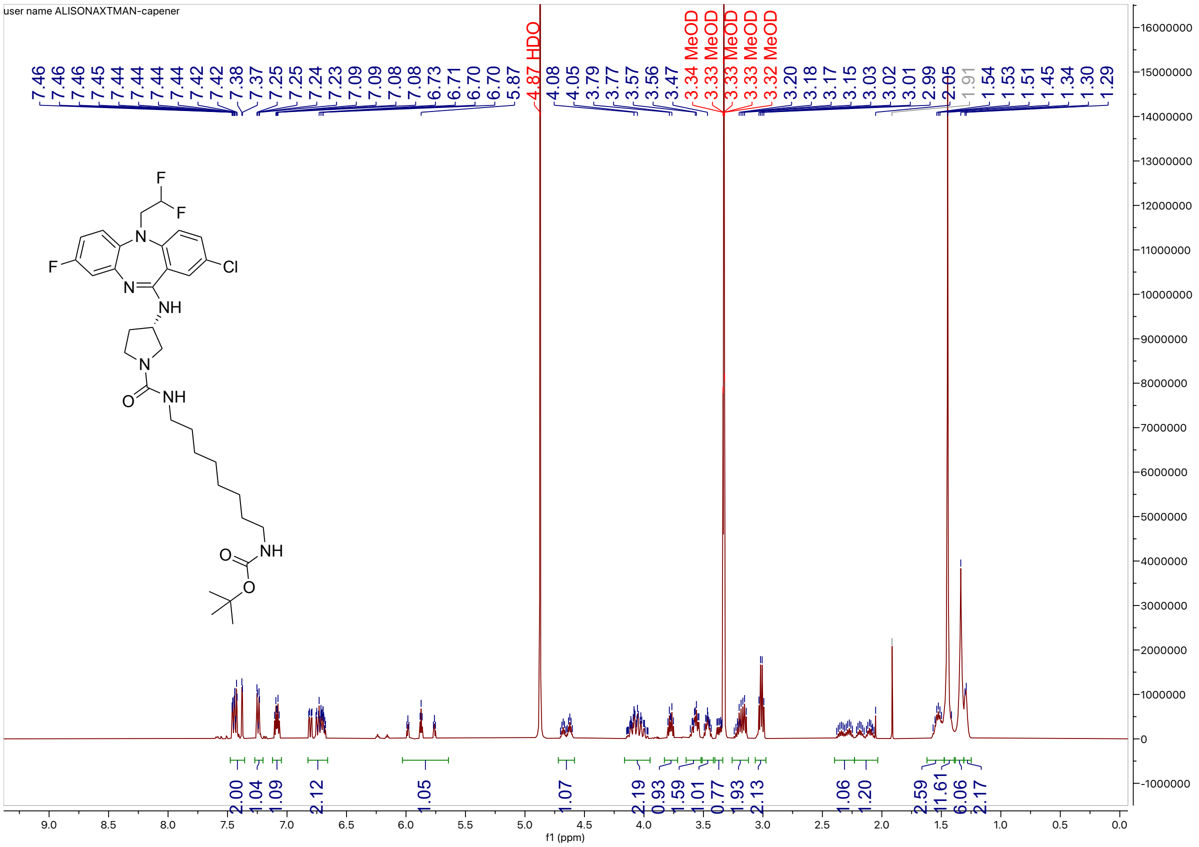

[1]H NMR (400 MHz, CD$_3$OD) for compound **4**

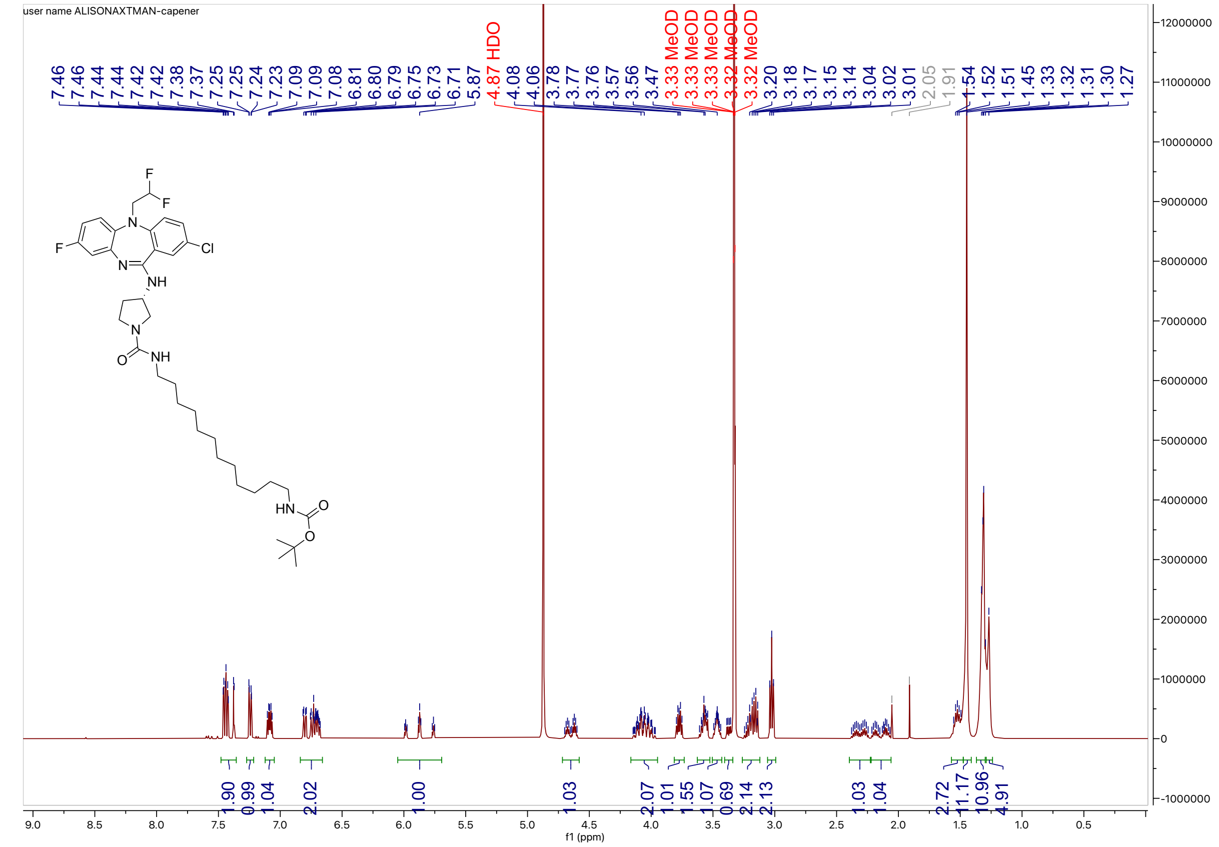


[1]H NMR (400 MHz, CD$_3$OD) for compound **5**

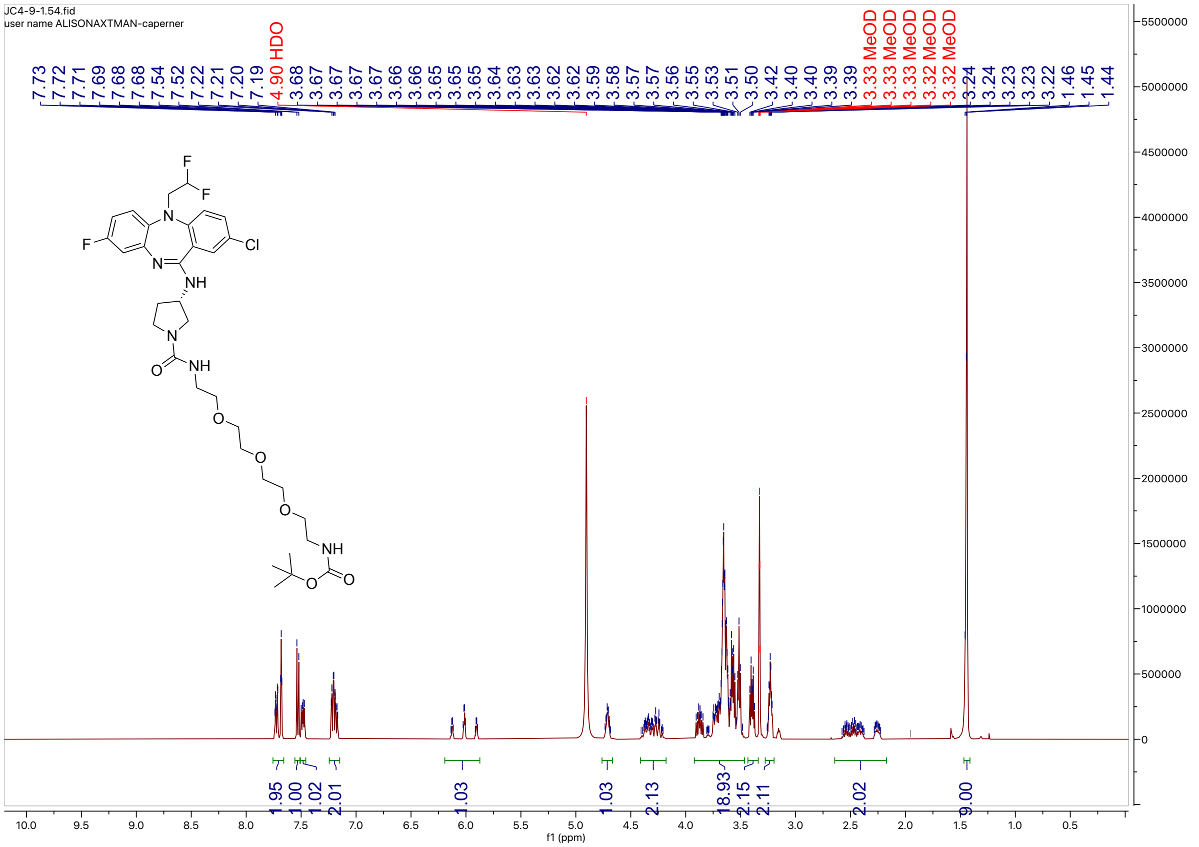

[1]H NMR (400 MHz, $CD_3OD$) for compound **6**

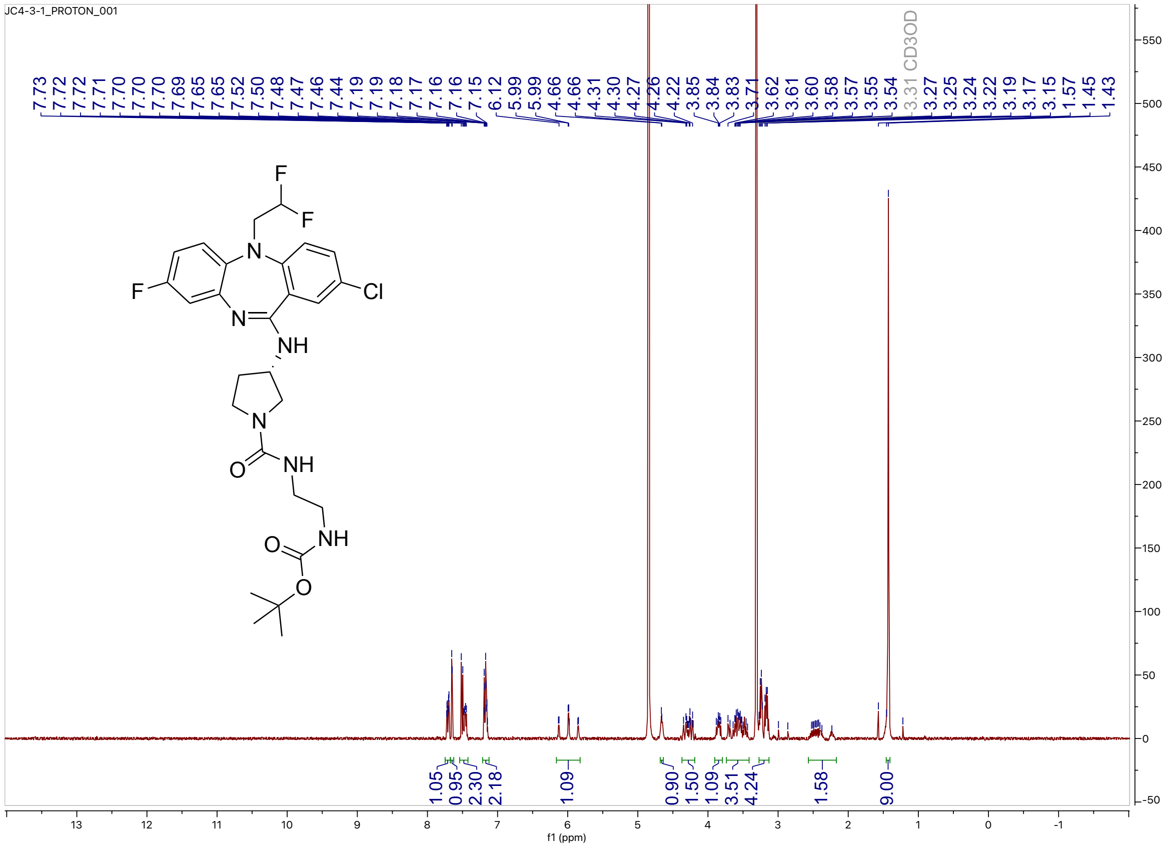

[1]H NMR (400 MHz, $CD_3OD$) for compound **7**

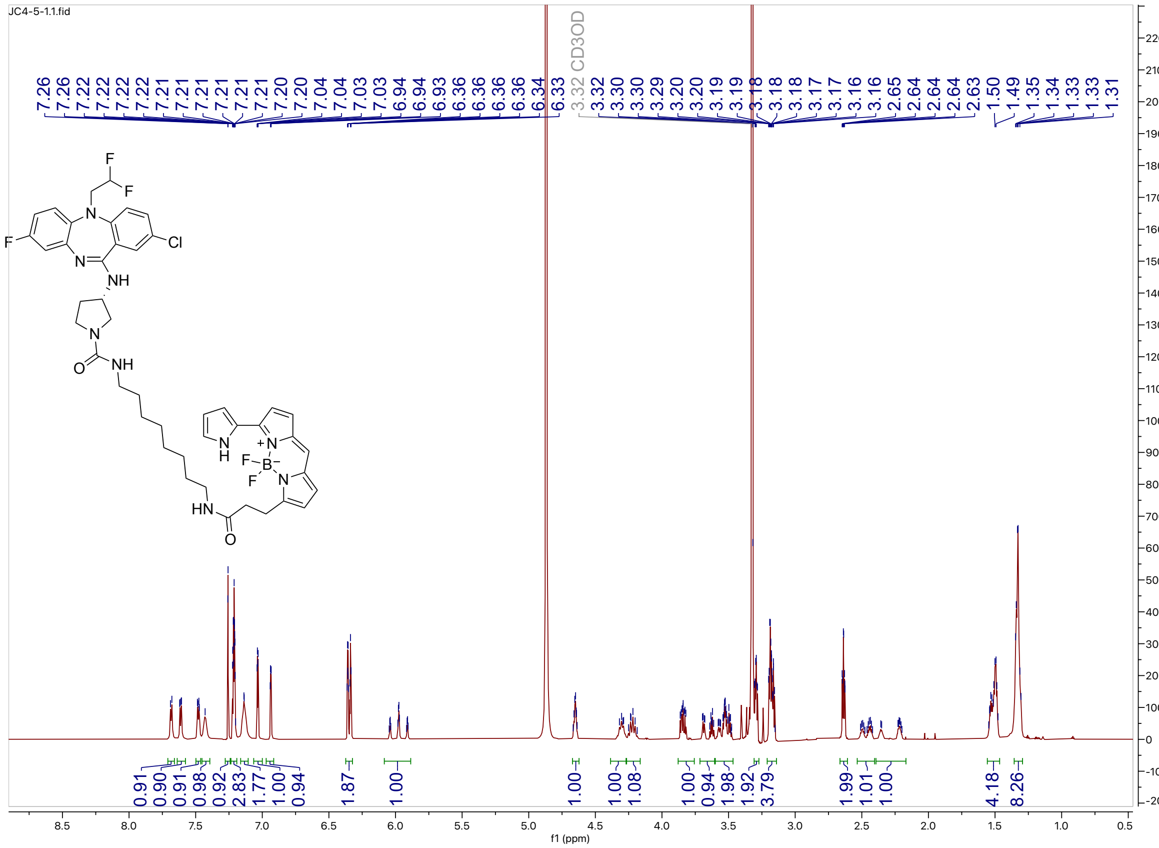

HPLC trace for compound 7

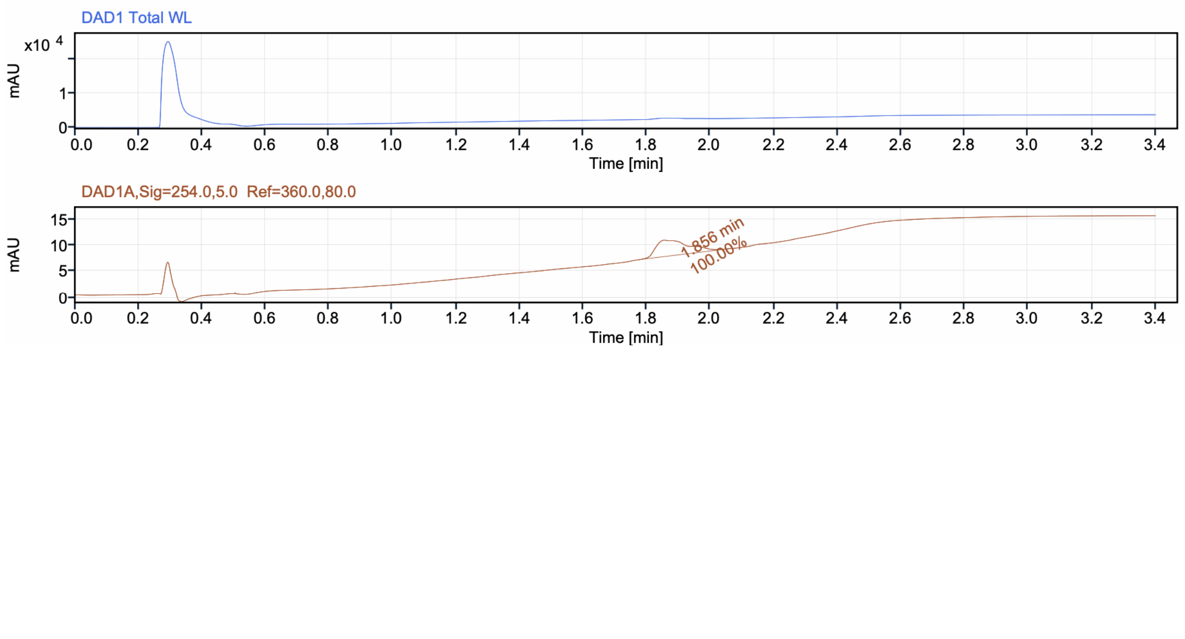

[1]H NMR (400 MHz, $CD_3OD$) for compound **8**

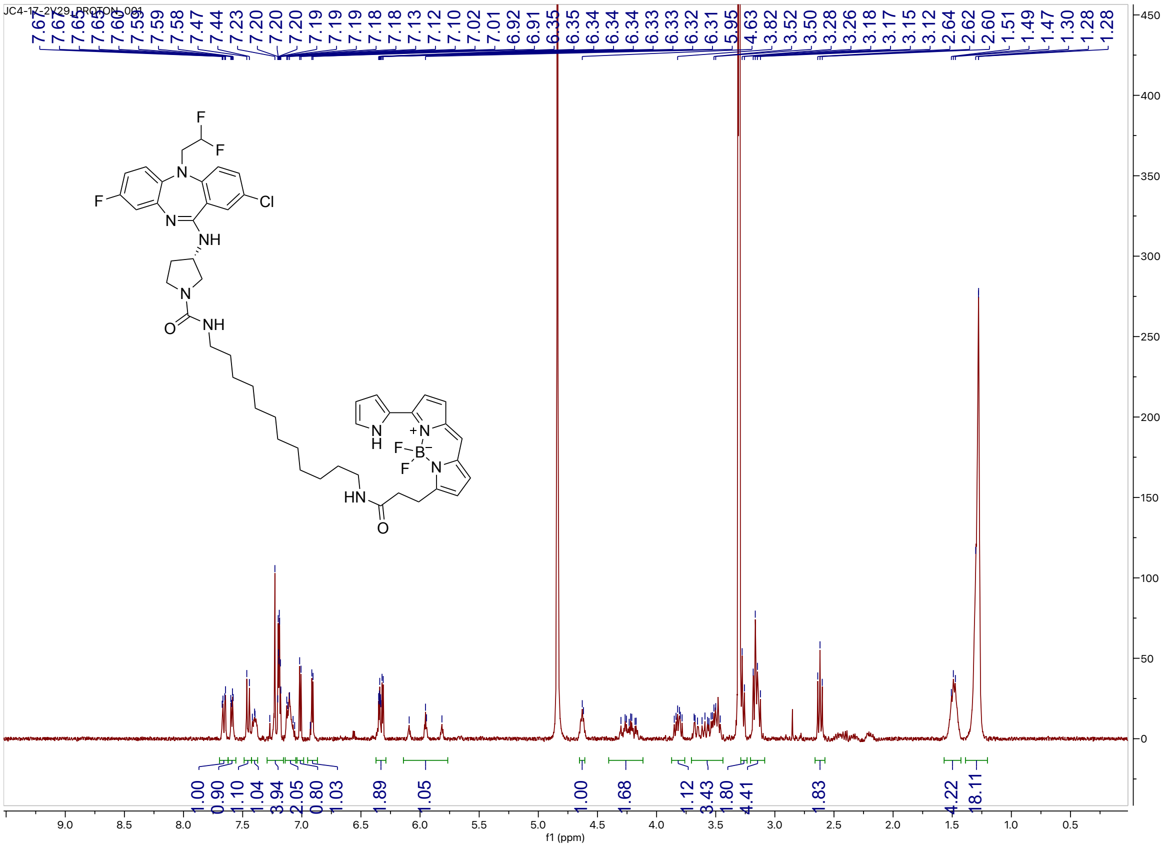

HPLC tracer for compound **8**

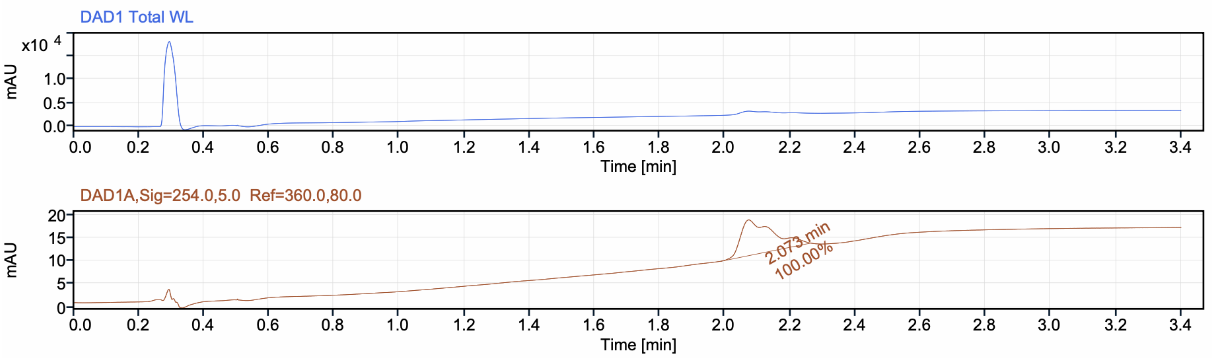

[1]H NMR (400 MHz, $CD_3OD$) for compound **9**

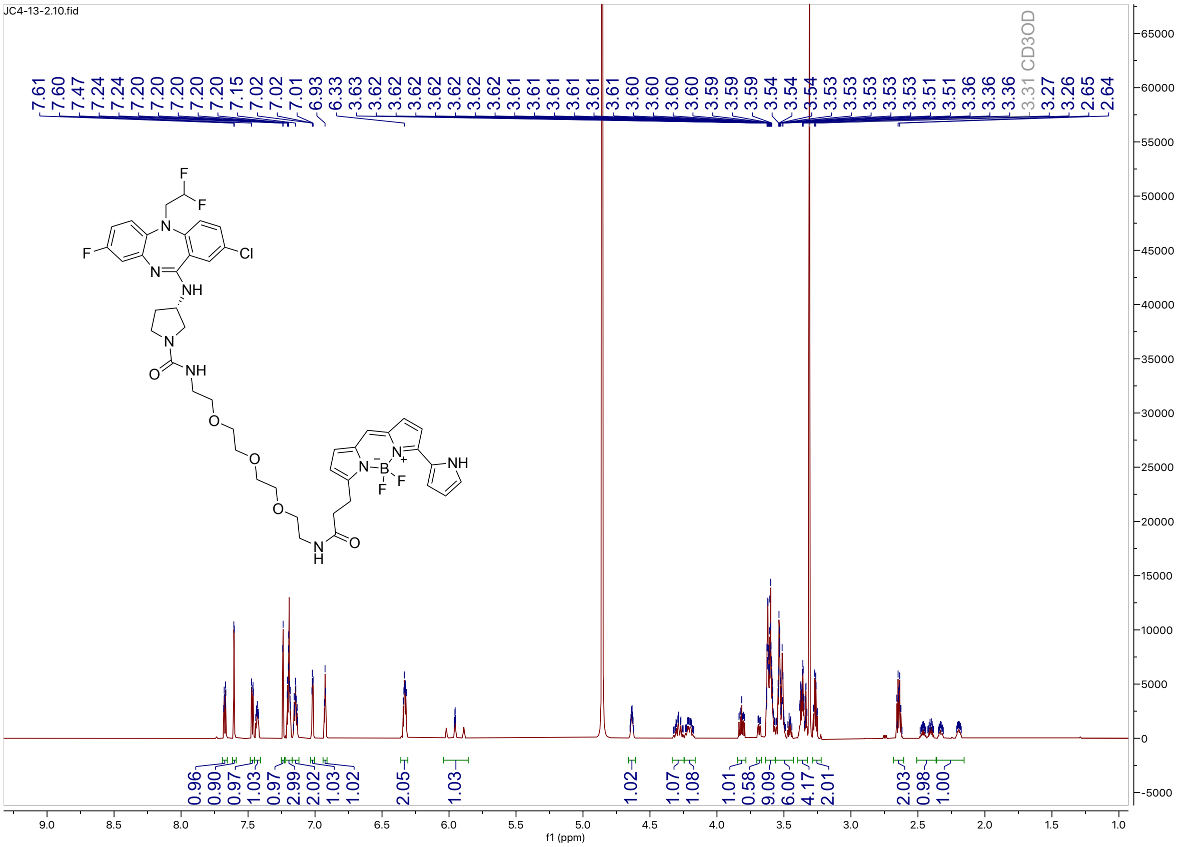

HPLC trace for compound **9**

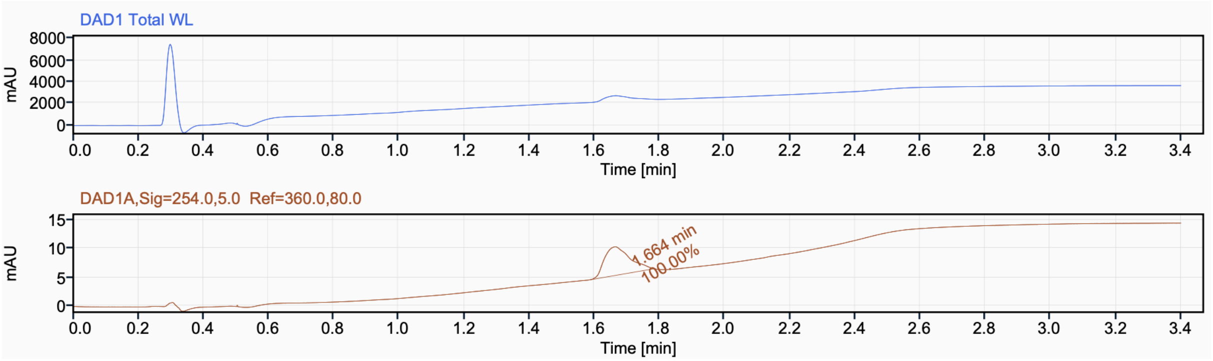

[1]H NMR (400 MHz, $CD_3OD$) for compound **10**

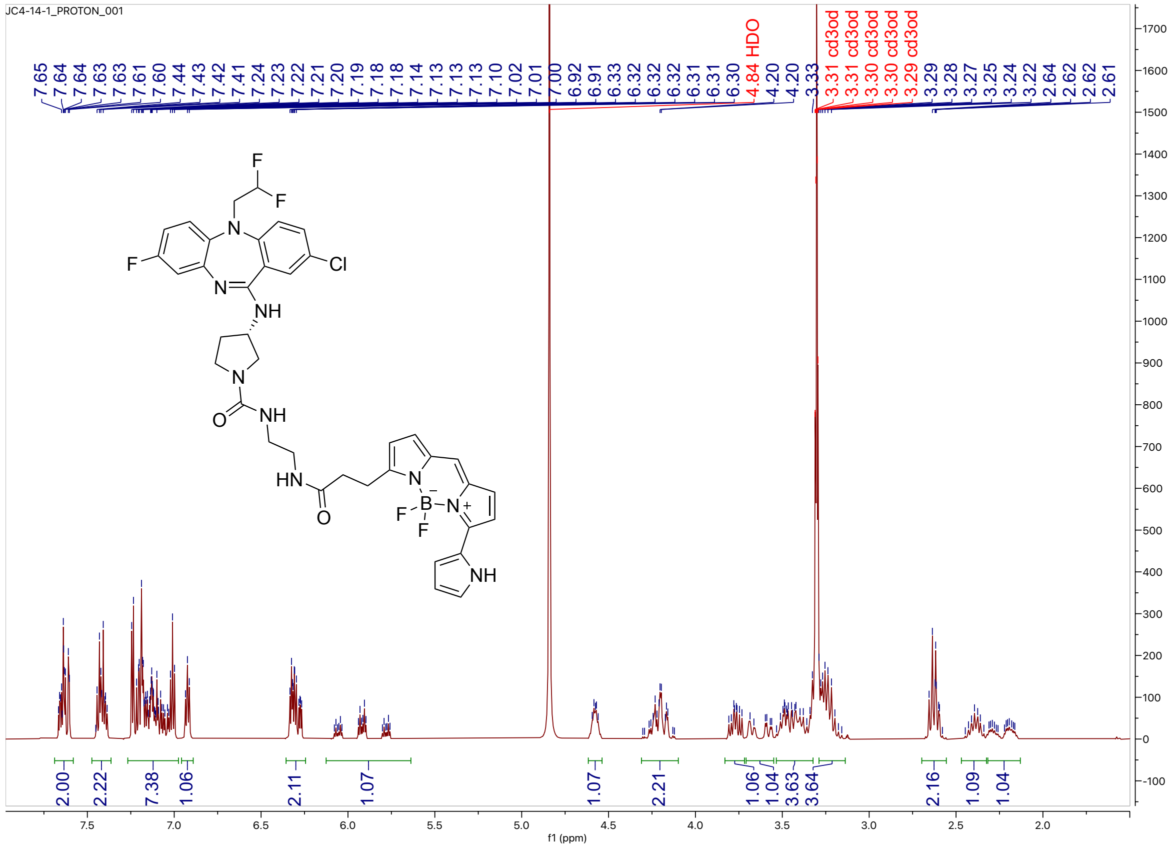

HPLC trace for compound **10**

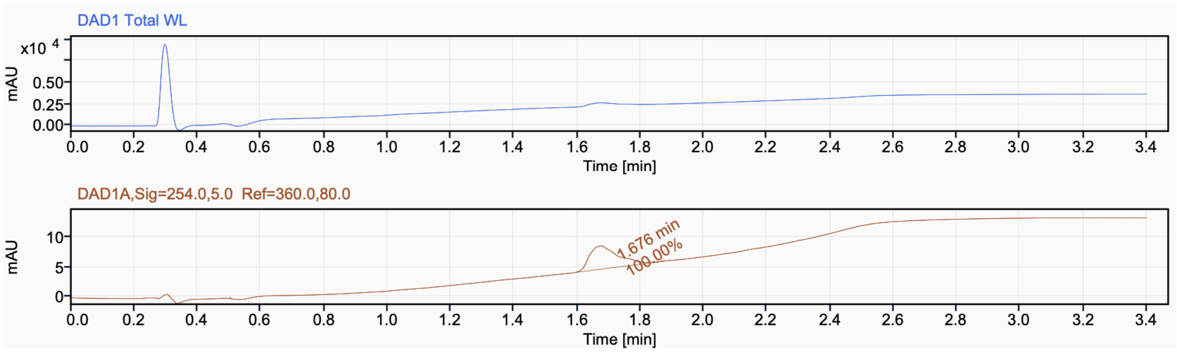
